# Ist2, a protein involved in phosphatidylserine transport, is an ER lipid scramblase

**DOI:** 10.1101/2025.02.13.638165

**Authors:** Heitor Gobbi Sebinelli, Camille Syska, Hafez Razmazma, Véronique Albanèse, Ana Rita Dias Araujo, Cecile Hilpert, Cédric Montigny, Christine Jaxel, Manuella Tchamba, Karolina Belingar, Juan Martín D’Ambrosio, Luca Monticelli, Guillaume Lenoir, Alenka Čopič

## Abstract

Lipid scramblases allow passive flip-flop of phospholipids between bilayer leaflets, thereby promoting membrane symmetry. At the endoplasmic reticulum (ER), where phospholipid synthesis is restricted to one of the two leaflets, scramblase activity should be essential for equilibrated membrane growth. However, phospholipid scramblases at the ER are poorly understood. The yeast protein Ist2 contains an ER domain and a cytosolic tail that binds the plasma membrane (PM) and participates in the transfer of phosphatidylserine (PS). Here, we show both *in vitro* and *in silico* that the ER- domain of Ist2, which bears homology to the TMEM16 proteins, possesses a lipid scramblase activity. Ist2 activity is not regulated by Ca^2+^, in contrast to TMEM16 proteins, but is affected by the lipid composition of the bilayer used in simulations. In cells, we do not find a strong impact of the scramblase domain of Ist2 in on PS distribution; however, its over-expression or deletion affects processes at the ER such as vesicular transport, lipid droplet biogenesis and general phospholipid transport, with a specific contribution of residues important for lipid scrambling. Our study therefore identifies the first dedicated phospholipid scramblase in yeast and demonstrates that membrane asymmetry can impact diverse membrane-remodeling processes at the ER.

## Introduction

A hallmark of eukaryotic cells is their highly diversified membrane system, which is essential for the organization and compartmentalization of cellular functions. Membrane diversity is reflected by large differences in their bulk lipid composition, driven by localized lipid synthesis and by the action of cytosolic lipid transfer proteins (LTPs), which mediate selective transport of lipids between different organelles (Holthuis and Menon, 2014; Lenoir et al., 2021). For example, phosphatidylserine (PS) and cholesterol are both enriched at the plasma membrane (PM) through the action of specific LTPs (Drin, 2023). Diversity in lipid composition can also be observed within the same membrane either as lateral inhomogeneity or as difference between the two leaflets (trans-bilayer asymmetry).

Membrane trans-bilayer asymmetry is most striking and well-studied in the case of the PM. The cytosolic leaflet of the PM is highly negatively charged due to an abundance of anionic phospholipids, such as PS, whereas the exoplasmic leaflet is almost devoid of these phospholipids but enriched in sphingolipids (Bretscher and Raff, 1975; Lorent et al., 2020). It is also well established that, in contrast to the PM, the cytosolic leaflet of the ER is poorly charged (Bigay and Antonny, 2013; Yeung et al., 2008). However, when we zoom in on the ER membrane, the data on its precise lipid organization is scarce and often conflicting. Taking again PS as an example, Fairn and coworkers have shown that it is largely confined to the luminal ER leaflet (Fairn et al., 2011). In contrast, a subsequent study suggests that PS concentration may be much more equilibrated between the two ER leaflets, or possibly even reversed (Tsuji et al., 2019). While much more data is clearly needed, these conflicting observations may reflect the highly dynamic and multifunctional nature of the ER, which hosts many processes such as protein and lipid synthesis, membrane transport and lipid transport via LTPs, and the biogenesis of organelles such as lipid droplets (LDs) and autophagosomes.

Three mechanisms are essential for the establishment of bilayer asymmetry: (1) localized phospholipid synthesis (Bell et al., 1981; Long et al., 2024; Vance et al., 1977); (2) slow rate of spontaneous phospholipid exchange (flip-flop) between the two leaflets (Bretscher and Raff, 1975), and (3), the action of phospholipid flippases, which use ATP hydrolysis to promote phospholipid flow from the extracellular/luminal towards the cytosolic membrane leaflet (Sakuragi and Nagata, 2023). In opposition to these mechanisms, phospholipid scramblases enable flow of phospholipids down their concentration gradient, disrupting bilayer asymmetry (Sebinelli et al., 2024). A well-studied example is the TMEM16 protein family, which includes several Ca^2+^- activated PM lipid scramblases (Brunner et al., 2014; Falzone et al., 2019; Kalienkova et al., 2021; Yang et al., 2012). Mutations in TMEM16F are associated with Scott Syndrome, a bleeding disorder resulting from defective PS exposure in activated platelets (Jan and Jan, 2024; Suzuki et al., 2010). An increase in cytosolic Ca^2+^ concentration induces a conformational change that leads to the opening of a hydrophilic lipid permeation pathway, enabling the passage of phospholipid headgroups via the so-called ‘credit-card’ model (Pomorski and Menon, 2006).

Because most phospholipids are synthesized in one or the other ER leaflet, scramblase activity at the ER would be essential for equilibrated membrane growth (Chauhan et al., 2016). However, specific ER-localized lipid scramblases only recently began to be identified. Two mammalian ER proteins of the DedA family, TMEM41B and VMP1, scramble phospholipids *in vitro* and the scramblase activity of TMEM41B has also been assessed in cells (Ghanbarpour et al., 2021; Huang et al., 2021; Li et al., 2021). Furthermore, TMEM16K, a TMEM16 family scramblase which localizes to the ER in mammalian cells, has also been shown to facilitate lipid scrambling in a Ca^2+^-dependent manner (Bushell et al., 2019; Tsuji et al., 2019). The activities of these proteins are often coupled to other lipid/membrane transport pathways (Hama et al., 2022). TMEM41B and VMP1 function in autophagy, where they may work in concert with the LTP ATG2 and the autophagosomal scramblase ATG9 to promote autophagosomal membrane expansion (Ghanbarpour et al., 2021; Matoba et al., 2020; Van Vliet et al., 2022). TMEM41B scramblase activity has been implicated in the production of lipoprotein particles and its deletion is often associated with accumulation of cytosolic LDs (Huang et al., 2021; Moretti et al., 2018; Morita et al., 2018; Wu et al., 2024). TMEM16K has been shown to affect endo-lysosomal transport pathway and localizes at ER-endosome contact sites, whereas TMEM16H functions at ER-PM contact sites (Jha et al., 2019; Petkovic et al., 2020). In addition to these recently identified specific ER scramblases, other ER proteins may moonlight as lipid scramblases. G-protein coupled receptors (GPCRs) have been proposed to function as constitutive scramblases in the ER while en route to their final destination at the PM, where they would be inhibited by cholesterol (Goren et al., 2014; Morra et al., 2022). Very recently, it has also been proposed that all ER membrane insertases involved in the synthesis of ER membrane proteins possess additional lipid scramblase activity (Li et al., 2024). Therefore, the field has recently witnessed a sharp turn from too few known ER scramblases to a host of different ER scramblases, raising the question of coordination of the various processes at the ER.

In budding yeast, a striking example of a protein machinery that could combine different lipid transport activities at the ER is provided by the Ist2-Osh6 complex. Ist2 localizes to ER-PM membrane contact sites and is composed of two distinctive parts: an ER-embedded transmembrane domain (TMD) that is preceded by a short cytosolic N-terminus (N-ter), and a largely disordered C-terminal (C-ter) tail that tethers the ER to the PM (Collado et al., 2019; Hoffmann et al., 2019; Kralt et al., 2014; Manford et al., 2012; Wolf et al., 2012). The cytosolic LTPs Osh6 and Osh7 (Filseck et al., 2015; Maeda et al., 2013) bind to the cytosolic tail of Ist2, enabling their localization and efficient PS transfer from the ER to the PM (D’Ambrosio et al., 2020; Wong et al., 2021). The TMD of Ist2 contains 10 putative transmembrane helices and shows sequence similarity with the TMEM16 protein family, with conservation of some residues previously identified as critical for scrambling (Fig 1A, B) (Brunner et al., 2014; Falzone et al., 2019; Kalienkova et al., 2019). This suggests that Ist2 could be a phospholipid scramblase in the ER, although an initial attempt using purified and reconstituted Ist2 failed to reveal any flip-flop activity for this protein (Malvezzi et al., 2013). In the present work, we investigate whether Ist2 is a phospholipid scramblase and how this activity at the ER might impact PS transfer and other ER functions.

**Figure 1.**
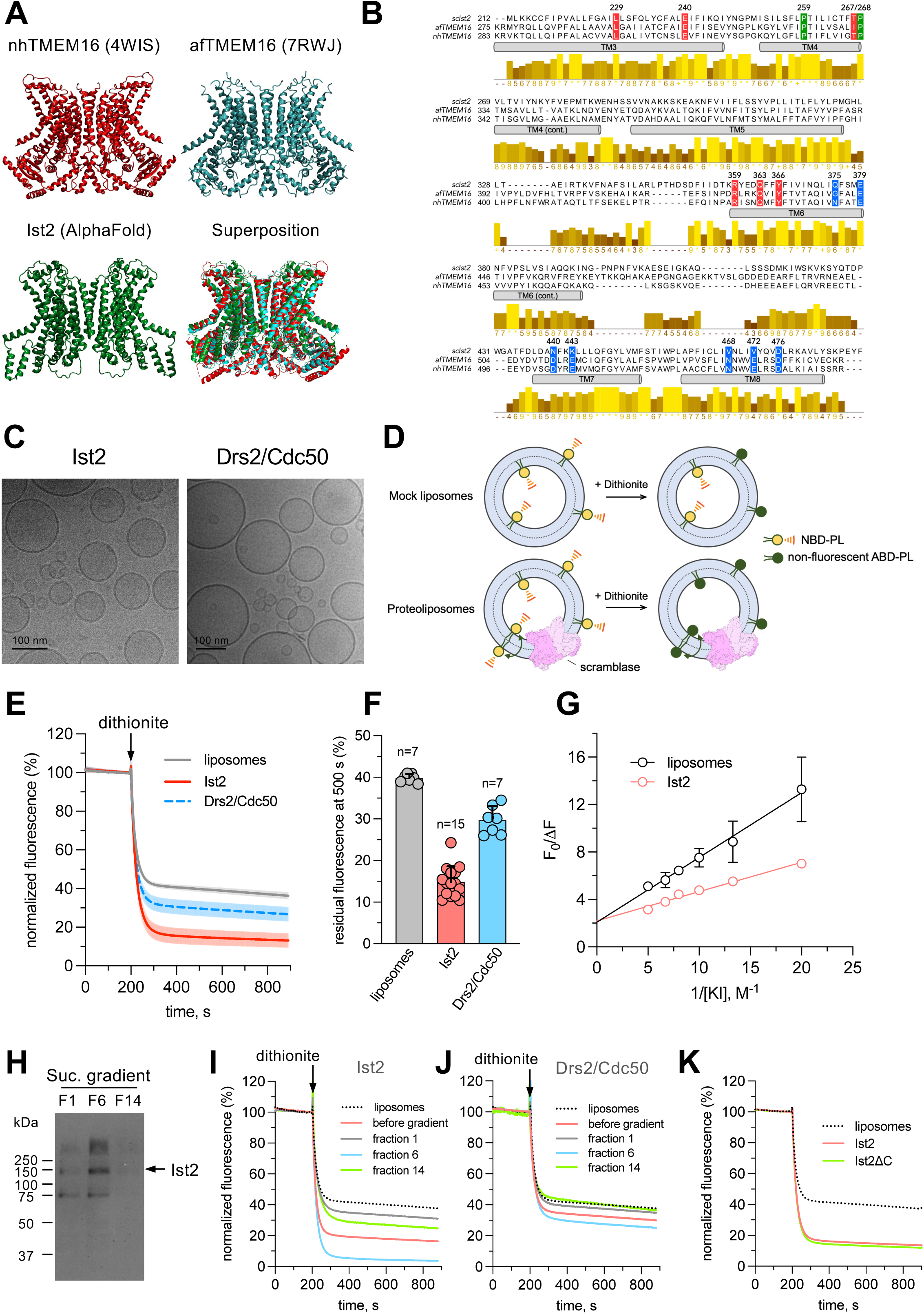
Reconstitution of Ist2 into proteoliposomes and dithionite scrambling assay. **(A)** Comparison of nhTMEM16 (4WIS) and afTMEM16 (6E1O) structures with that of Ist2 predicted by AlphaFold2. **(B)** Sequence alignment of Ist2 from *S. cerevisiae* with *N. haematococca* TMEM16 (nhTMEM16) and *A. fumigatus* TMEM16 (afTMEM16) (see also phylogenetic tree in Fig. S1A). Sequences were aligned using the MUSCLE server (EMBL-EBI). The predicted transmembrane helices (TM) of Ist2 based on its AlphaFold2 model are shown as grey cylinders. The residues critical for phospholipid scrambling in nhTMEM16 and the two prolines in TM4 serving as pivots for helix rearrangements, as well as the corresponding residues in afTMEM16 and Ist2, are highlighted in red and green, respectively (Jiang et al 2017; Lee et al, 2018; Kalienkova et la, 2019). Residues defining the Ca^2+^-binding sites in nhTMEM16 are highlighted in blue (Brunner et al, 2014). Conservation is shown below the sequences (brown – yellow = least – most conserved). Uniprot identifiers: Ist2 (P38250), nhTMEM16 (C7Z7K1), afTMEM16 (Q4WA18). **(C)** Cryo-electron micrographs of PLs containing Ist2 or the Drs2/Cdc50 lipid flippase complex. The average diameter is ∼ 80-100 nm for both PLs Scale bar = 100 nm. **(D)** Schematic for the dithionite-based lipid transport assay. Unilamellar vesicles are formed in the presence of a trace amount of fluorescent 7-nitro-2,1,3-benzoxadiazol-4-labeled phospholipid (NBD-phospholipid), which can be reduced by dithionite to a non-fluorescent 7-amino analog. In the case of lipid flip-flop, dithionite treatment of scramblase-reconstituted PLs should result in 100% fluorescence loss, whereas treatment of mock liposomes should result in 50% fluorescence loss as only NBD in the outer leaflet is available to react with dithionite. **(E)** NBD-PS scrambling assay. C12-NBD-PS-containing liposomes or PLs were stirred in a cuvette. The initial fluorescence was recorded for 200 s before dithionite addition. The shaded area around each trace represents the standard deviation for each fluorescent point. **(F)** Quantification of residual fluorescence in liposomes versus PLs measured 500s after dithionite addition (panel E). Data are a mean ± s.d. of 7 to 15 biological replicates (independent reconstitutions). **(G)** Collisional quenching of C12-NBD-PS fluorescence with iodide ions in liposomes and Ist2-containing PLs. The data are as modified Stern-Volmer plots (see Materials and Methods). F_0_ is the fluorescence intensity of the sample in the absence of quencher, whereas F is the fluorescence intensity at a given iodide ion concentration. The fluorescence data points were fitted to a linear regression. The inverse of the y-intercept represents the fraction of C12-NBD-PS that is accessible to the quencher. Data are a mean ± s.d. of 3 biological replicates (independent reconstitutions) for both liposomes and Ist2-containing PLs. Some of the error bars are not visible because they are smaller than the symbol. The equation of the linear regression for liposomes is Y=0.5462X + 2.074, meaning that 48% of NBD-lipids are accessible to the quencher, whereas for Ist2-containing proteoliposomes, the equation is Y=0.2469X + 2.191, meaning that 46% of NBD-lipids are accessible to the quencher. **(H)** Western-blot analysis of fractions collected upon fractionation of Ist2-mCherry-containing PLs on a sucrose gradient. **(I)** NBD-PS quenching of fractions recovered upon fractionation of Ist2 PL on a sucrose gradient, compared to Ist2 PL and liposomes before fractionation. **(J)** NBD-PS quenching of fractions recovered upon fractionation of Drs2/Cdc50 PL on a sucrose gradient, compared to PLs and liposomes before fractionation. **(K)** NBD-PS quenching of liposomes or proteoliposomes reconstituted with either full-length Ist2 or Ist2-ΔC.

We show using biochemical reconstitution and molecular dynamics (MD) simulations in an ER-like bilayer that Ist2 can scramble phospholipids in a non-selective and Ca^2+^-independent manner. In yeast, deletion of the ER domain of Ist2 does not have a strong impact on cellular phospholipid levels or their steady-state distribution. However, we find that deleting this domain is lethal in the context of the PS-synthesis mutant *cho1*Δ. We further show that the TMD of Ist2 affects the ER vesicular export pathway as well as LD abundance and composition, and that these phenotypes are rescued by mutations in conserved residues predicted to be critical for phospholipid scrambling. Overall, our results point to the inter-connectedness of ER functions and underline the importance of the regulation of ER trans-bilayer lipid distribution.

## Results

### Ist2 functions as a phospholipid scramblase *in vitro*

To get insights into the function of the ER-embedded TMD of Ist2, we first performed structural homology modelling and sequence alignment. For homology modelling, we used the portion of the *Saccharomyces cerevisiae* Ist2 sequence homologous to TMEM16 proteins, i.e. from residue 1 to 600 (Ist2ΔC), excluding the disordered C-tail, which is unique to yeasts (D’Ambrosio et al., 2020). Comparison of Ist2ΔC with the top ten of homologous proteins identified by I-TASSER revealed that the two closest Ist2 orthologues belong to the TMEM16 family, whose structures have been experimentally determined, nhTMEM16 from *Nectria haematococca* (Brunner et al., 2014; Kalienkova et al., 2019) and afTMEM16 from *Aspergillus fumigatus* (Falzone et al., 2019) (Fig. S1A). Both nhTMEM16 and afTMEM16, as well as other TMEM16 family proteins, form homodimers, suggesting that Ist2 TMD also dimerizes (Fig. 1A). These proteins have been shown to function as Ca^2+^-activated phospholipid scramblases and/or as Ca^2+^-activated chloride channels (Brunner et al., 2014; Falzone et al., 2019; Jiang et al., 2017; Malvezzi et al., 2013) . Sequence alignment shows a strong conservation of residues critical for lipid scrambling between nhTMEM16, afTMEM16 and Ist2, whereas residues involved in Ca^2+^-binding are not all conserved in Ist2 (Fig. 1B).

To test whether Ist2 scrambles phospholipids, we used a well-established fluorescence-based dithionite assay (Ploier and Menon, 2016). We first purified Ist2 fused to a biotin-acceptor domain following its overexpression in budding yeast, using a previously established procedure (Azouaoui et al., 2014; Dieudonné et al., 2022). In addition, mCherry was fused to Ist2 to facilitate its detection. Both full-length Ist2 and Ist2ΔC could be efficiently purified upon extraction with n-dodecyl-β-D-maltoside (DDM) from yeast membranes (Fig. S1B,C). Next, Ist2 was reconstituted into proteoliposomes (PLs) containing a small percentage of nitrobenzoxadiazole (NBD)-labeled lipid. Despite some protein loss during the reconstitution process, Ist2 was effectively recovered after detergent removal with Bio-beads (Fig. S1D) and unilamellar PLs were generated, as assessed by cryo-electron microscopy (Fig. 1C). The average diameter deduced from the observation of cryo-electron micrographs was ∼80 nm for Ist2 PLs and 100 nm for control liposomes.

In the dithionite assay, the emission of the NBD-labeled lipid is irreversibly quenched due to the reduction of NBD into a non-fluorescent ABD moiety by dithionite (Fig. 1D). In control liposomes, 50% of the initial fluorescence is expected to be bleached as only NBD-lipids in the outer leaflet are accessible, whereas in PLs containing a scramblase that exchanges lipids from one leaflet to the other, all NBD lipids should be accessible and fluorescence reduction should in principle be complete. As shown in Fig. 1E and 1F, dithionite addition to Ist2-containing PLs induced a significantly larger fluorescence reduction than for control liposomes. Fluorescence reduction by dithionite of protein-free liposomes is higher than the predicted value (∼60% vs ∼50%); however, this is consistent with the relatively small size of the PLs. When a control, ATP-dependent Drs2/Cdc50 lipid flippase complex was reconstituted in PLs, fluorescence reduction in the absence of ATP was much less pronounced than for Ist2 PLs (Fig. 1E,F). We also confirmed that the presence of an mCherry tag on Ist2 did not affect fluorescence reduction by dithionite (Fig. S1E).

To exclude the possibility that the increased fluorescence reduction upon addition of dithionite is due to the preferential incorporation of NBD-PS in the outer leaflet of Ist2 PLs, we used iodide collisional quenching of the NBD fluorophore (Goren et al., 2014). In this assay, membrane-impermeable iodide ions reversibly quench the fluorescence of the NBD-PS located in the outer leaflet of the proteoliposomes, reporting on its steady-state distribution. Since scramblases continuously exchange lipids from one leaflet of the PL to the other, iodide treatment should result in 50% quenching. Analysis of the quenching experiment revealed that NBD-PS is symmetrically distributed in both liposomes and proteoliposomes (Fig. 1G). Furthermore, the observed 50% protection of NBD-PS demonstrates that Ist2 PLs are tight to iodide ions. Further purification of PLs on a sucrose gradient confirmed the successful reconstitution of Ist2 into vesicles, as Ist2 co-sedimented with NBD-lipids (Fig. 1H, S1F). The top fraction (F1) contained pure lipid vesicles whereas the fluorescent ring observed in a lower fraction (F6) mostly corresponded to PLs. Consistent with this fractionation, fluorescence reduction was almost complete when dithionite was added to the PL-enriched fraction (F6, Fig. 1I). In contrast, little change was observed for the Drs2/Cdc50 control sample (Fig. 1J). We also asked whether the disordered C-ter tail of Ist2 contributes to lipid scrambling mediated by Ist2. Ist2-ΔC reconstituted similarly to full-length Ist2, and the truncation of the C-ter tail did not alter the transbilayer movement of NBD-PS (Fig. 1K, Fig. S1G). Altogether, these data indicate that the TMD of Ist2 promotes lipid scrambling after its purification and reconstitution in proteoliposomes.

### Ist2 scrambles phospholipids in a Ca^2+^-independent manner with some selectivity

The TMEM16 proteins characterized so far have been shown to be stimulated by Ca^2+^. Upon Ca^2+^ binding, a lateral groove called the subunit cavity opens, thereby providing a pathway large enough for lipids to cross the membrane (Falzone et al., 2019; Kalienkova et al., 2019). In contrast to other TMEM16 members, Ist2 was essentially insensitive to Ca^2+^, as the addition of Ca^2+^ to reconstituted Ist2 did not alter its ability to transport NBD-PS (Fig. 2A,B). Guided by the homology between Ist2 and proteins of the TMEM16 superfamily (Fig. 1A,B), we built models for TMD of Ist2 using AlphaFold2 (Jumper et al., 2021) and trRosetta (Du et al., 2021). AlphaFold provided model structures of homodimers very similar to the nhTMEM16 X-ray structure (PDB ID: 4WIS) (Brunner et al., 2014), with an open subunit cavity; trRosetta, instead, provided also model structures similar to the conformation of nhTMEM16 in a closed state (PDB ID: 6QMB) (Kalienkova et al., 2019) (Fig. 1A). Two Ca^2+^ ions were initially inserted in both the open and closed structural model, then the proteins were embedded in lipid bilayer membranes and we performed MD simulations at the all-atom level to assess the stability of calcium binding. In all simulations, in the absence of restraints, Ca^2+^ ions dissociated spontaneously and very rapidly from the protein, suggesting that calcium binding is weak (Fig. 2C). This finding is consistent with our *in vitro* measurements and with the observation that most of TMEM16 residues involved in calcium binding are not conserved in Ist2 (Fig. 1B and Fig. 2A,B).

**Figure 2.**
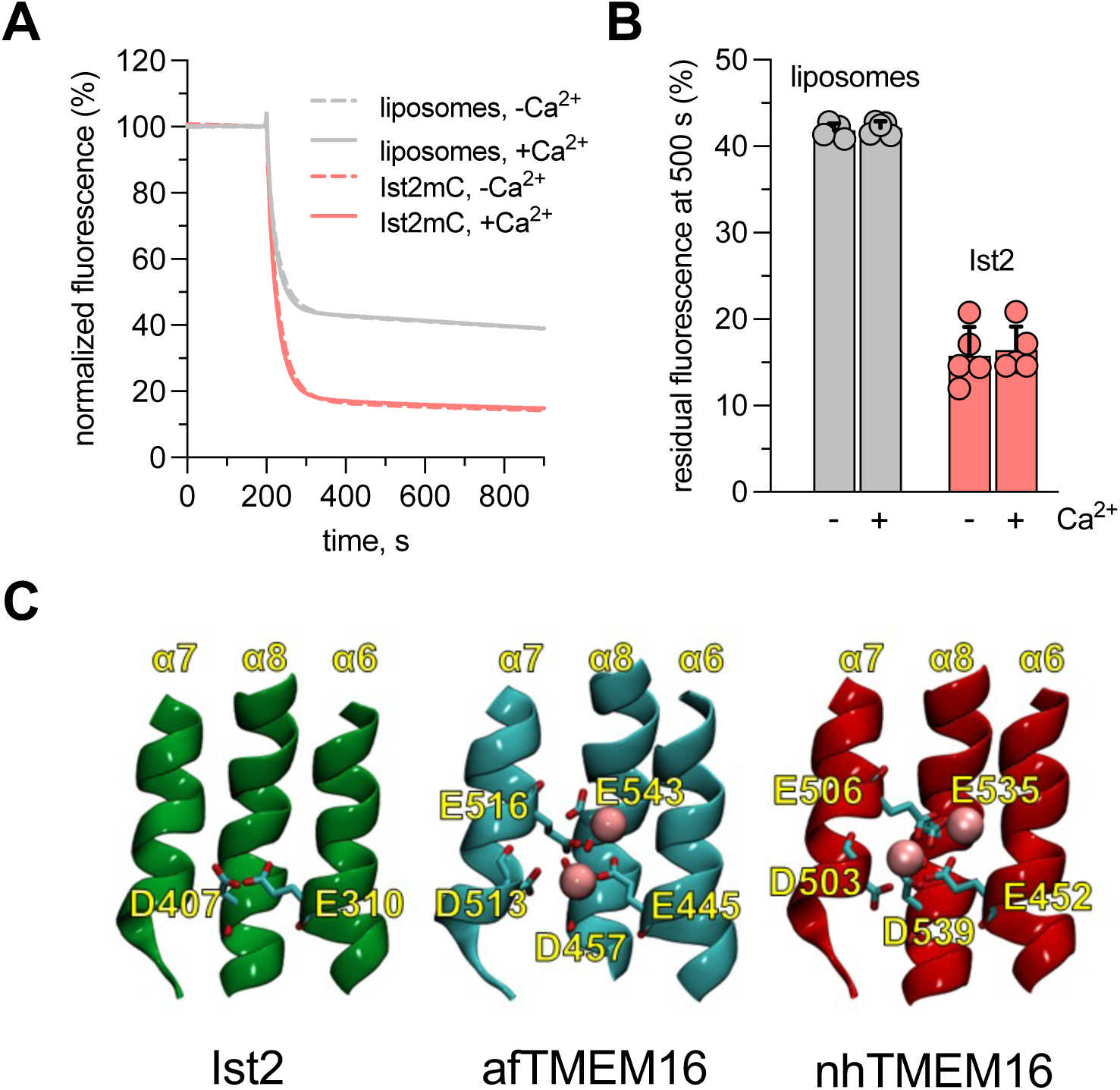
Ist2 scrambling activity is not sensitive to Ca^2+^. **(A)** Effect of Ca^2+^ on Ist2-mediated NBD-lipid scrambling. The effect of Ca^2+^ was assessed by adding x mM Ca^2+^ to both liposomes and Ist2-containing PLs. **(B)** Quantification of residual fluorescence in liposomes versus PLs upon dithionite quenching. The residual fluorescence measured in panel A at 500 s was plotted. Data are a mean ± s.d. of 5 biological replicates (independent reconstitutions). **(C)** Comparison of the region involved in Ca^2+^ binding between Ist2, afTMEM16, and nhTMEM16. (Only 2 out of 5 negatively charged residues involved in Ca2+ binding in TMEM16 are conserved in Ist2 – see Fig. 1B)

All-atom simulations were used to fine-tune coarse-grained (CG) models for the protein, using both the open and closed conformation; we did not include Ca^2+^ in the CG simulations. For each conformation we performed multiple long time-scale (20 µs) CG MD simulations using both a simple lipid mixture and a more complex one, mimicking the composition of the ER membrane (Reinhard et al., 2024) (see Table S3 for the lipid composition used in MD simulations, Table S4 for a full list of simulations, and the Methods section for the details of the simulation setup). Simulations provided a clear and reproducible result: a high number of flip-flop events were recorded in simulations of the open Ist2 conformation, and close to none for the closed conformation, irrespective of the membrane composition (Fig. 3A, Table 1-4 and Supplemental Movie 1). The rate of phospholipid flip-flop was in the same range as what has been observed for other TMEM16 scramblases *in silico* (Stephens et al., 2025). By contrast, flip-flop of ergosterol, which does not require scramblases, was fast and not affected by Ist2 conformation (Table 2, 4). In both simple and complex ER-like membranes, we observed selectivity for different phospholipids, with a notable preference for PC, PS, and the following over-all ranking: PC>PS>PE>PI>PA (Table 1-2). Interestingly, the probability of flip-flop also depended on acyl chains, with shorter chains (16:1) promoting flip-flop (Table 2). The rate of phospholipid flip-flop was higher in simple compared to complex lipid mixture, which could be explained by the combined acyl chain and headgroup preference, as well as some increase in the thickness of the complex bilayer. However, no phospholipid preference was observed in the *in vitro* dithionite assay, in which NBD-PS, NBD-PC and NBD-PE where consistently transported at a similar rate (Fig. 3B,C).

**Figure 3.**
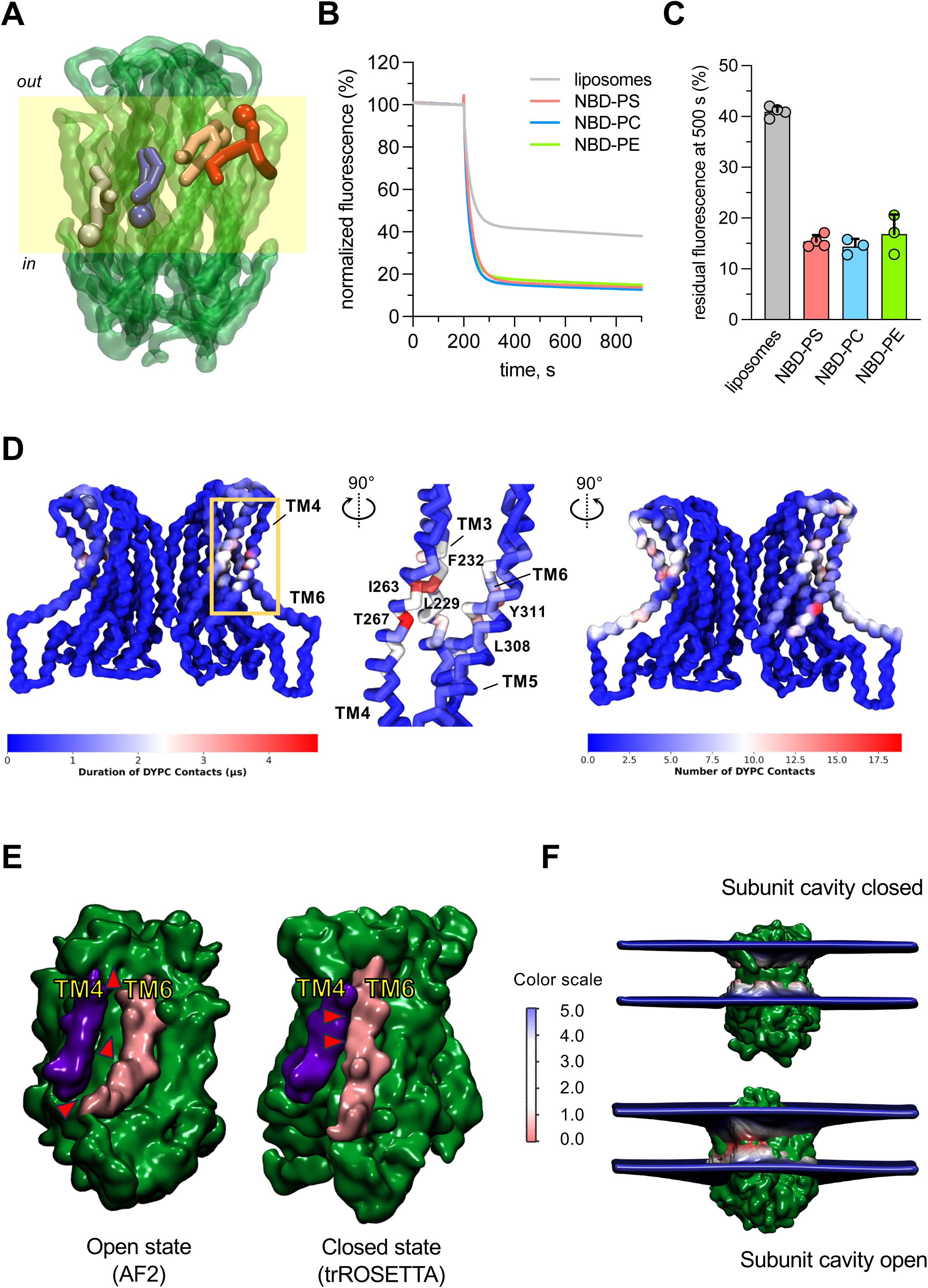
Mechanistic analysis of phospholipid scrambling by Ist2. **(A)** Illustration of the mechanism of scrambling of a PS lipid by Ist2. The protein is shown in green, the position of the lipid membrane is shaded in yellow, and one lipid is shown in 4 different time frames, corresponding to 4 different positions along the scrambling path. The lipid is shown with a thick stick representation and the serine head group is represented as a larger sphere, to highlight the flip-flop about half-way through the scrambling path. For a dynamic representation see the movie in supporting information. **(B)** Ist2 also scrambles NBD-PC and NBD-PE. Traces represent the mean of 3 to 5 transport experiments from independent reconstitutions. **(C)** Quantification of residual fluorescence in liposomes versus proteoliposomes upon dithionite quenching. The residual fluorescence measured in panel B at 500 s was plotted. Data are a mean ± s.d. of 3-5 biological replicates (independent reconstitutions). **(D)** The path followed by the lipid head groups during scrambling, predicted on the basis of lipid-protein contact during unbiased CG MD simulations. In the left panel, amino acids are colored based on the average duration of contacts with the PC head group; on the right panel, the color code indicates the average number of contacts during the MD simulation. At the center, the protein is rotated by 90 degrees and only 4 helices are shown, with residues showing the highest number of contacts highlighted. **(E)** Comparison of predicted models of Ist2 generated using AlphaFold 2 and trROSETTA. Note the variable proximity between TM4 (dark blue) and TM6 (pink) in the open and the close Ist2 model. **(F)** Membrane thickness, averaged over all simulations of the closed structural model (upper panel) and open structural model (lower panel). Thickness was calculated as the average distance between lipid head groups in each leaflet. The color scale indicates the average value of membrane in each position around the protein. Membrane thinning is visible around the open model, particularly in the position corresponding to the scrambling path.

**Table 1.** Scramblase activity of Ist2 (open state) in the simple lipid mixture.

| Lipid composition | | Total events / simulation (20 μs) | | | | Ist2 activity | |
| --- | --- | --- | --- | --- | --- | --- | --- |
| Lipid type<br>(1200 lipids in total) | Percentage | R1 | R2 | R3 | average ± SD | Flip-flops per ms per Ist2 dimer | Phospholipid preference index<sup>(1)</sup> |
| DYPC | 50% | 67 | 39 | 39 | 48 ± 16 | 2417 ± 808 | 1.3 |
| DYPE | 25% | 13 | 6 | 6 | 8 ± 4 | 417 ± 202 | 0.5 |
| DYPS | 25% | 21 | 12 | 12 | 15 ± 5 | 750 ± 260 | 0.8 |
| Phospholipids | 100% | 101 | 57 | 57 | 72 ± 25 | 3583 ± 1270 | 1.0 |

<sup>(1)</sup> Calculated by multiplying the average # of flip-flops with % of phospholipid species, divided by total # of flip-flop events.

**Table 2.** Scramblase activity of Ist2 (open state) in the complex lipid mixture.

| Lipid composition | | Total events / simulation (20 μs) | | | | Ist2 activity | |
| --- | --- | --- | --- | --- | --- | --- | --- |
| Lipid type<br>(1200 lipids in total) | Percentage<br>(PL only) | R1 | R2 | R3 | average ± SD | Flip-flops per ms per Ist2 dimer | Phospholipid preference index<sup>(1)</sup> |
| DYPC | 21% | 17 | 18 | 10 | 15 ± 4.4 | 750 ± 218 | 2.3 |
| YOPC | 21% | 9 | 9 | 3 | 7.0 ± 3.5 | 350 ± 173 | 1.1 |
| DYPE | 11% | 4 | 1 | 2 | 2.3 ± 1.5 | 117 ± 76 | 0.7 |
| YOPE | 11% | 2 | 0 | 2 | 1.3 ± 1.2 | 67 ± 58 | 0.4 |
| YOPS | 11% | 5 | 3 | 3 | 3.7 ± 1.2 | 183 ± 58 | 1.1 |
| POPI | 21% | 3 | 1 | 1 | 1.7 ± 1.2 | 83 ± 58 | 0.25 |
| YOPA | 5% | 1 | 0 | 0 | 0.3 ± 0.6 | 16 ± 29 | 0.2 |
| Phospholipids | 100% | 41 | 32 | 21 | 31 ± 10 | 1567 ± 501 | 1.0 |
| ERGO | 5% | 34704 | 34878 | 35062 | 34 881 ± 180 | 1 744 070 ± 8950 | n/a |

<sup>(1)</sup> Calculated by multiplying the average # of flip-flops with % of phospholipid species, divided by total # of flip-flop events.

**Table 3.**
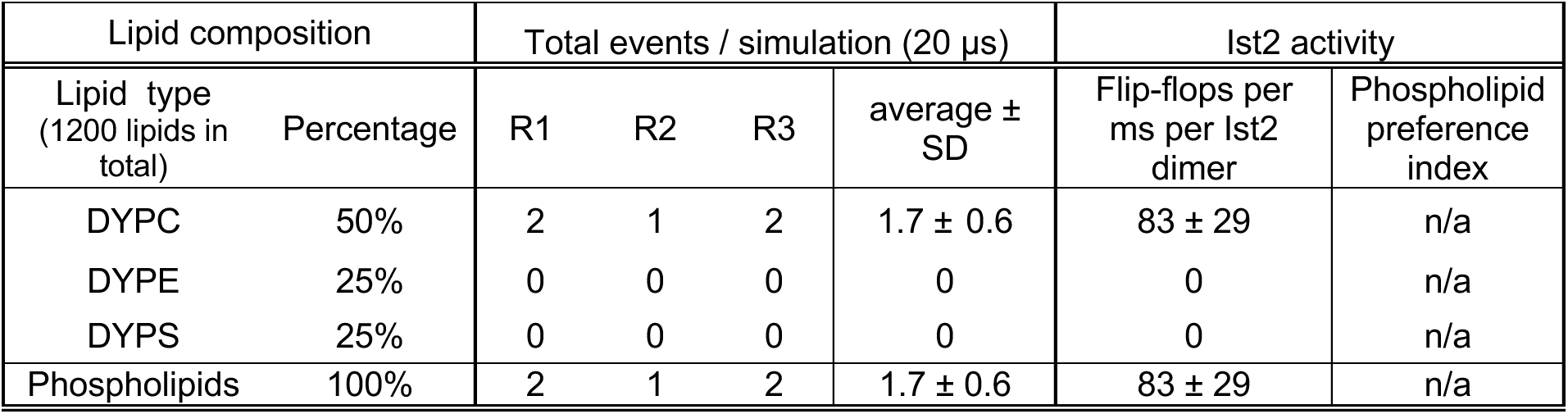
Scramblase activity of Ist2 TMD (closed state) in the simple lipid mixture.

**Table 4.** Scramblase activity of Ist2 TMD (closed state) in the complex lipid mixture.

| Lipid composition | | Total events / simulation (20 μs) | | | | Ist2 activity | |
| --- | --- | --- | --- | --- | --- | --- | --- |
| Lipid type<br>(1200 lipids in total) | Percentage<br>(PL only) | R1 | R2 | R3 | average ±<br>SD | Flip-flops per<br>ms per Ist2<br>dimer | Phospholipid<br>preference<br>index |
| DYPC | 21% | 0 | 0 | 1 | 0.3 ± 0.6 | 17 ± 29 | n/a |
| YOPC | 21% | 0 | 0 | 0 | 0 | 0 | n/a |
| DYPE | 11% | 0 | 0 | 0 | 0 | 0 | n/a |
| YOPE | 11% | 0 | 0 | 0 | 0 | 0 | n/a |
| YOPS | 11% | 0 | 0 | 0 | 0 | 0 | n/a |
| POPI | 21% | 0 | 0 | 0 | 0 | 0 | n/a |
| YOPA | 5% | 0 | 0 | 0 | 0 | 0 | n/a |
| Phospholipids | 100% | 0 | 0 | 0 | 0 | 0 | n/a |
| ERGO | 5% | 33156 | 33493 | 33269 | 33306 ± 172 | 1 665 300 ± 8576 | n/a |

We next identified the pathway for lipid transport by calculating protein-lipid contacts during the simulations. In all of the flip-flop events, lipids passed through the subunit cavity (between TM3- 6) facing the lipid bilayer on both sides of the dimer, as previously observed for other TMEM16 proteins (Fig. 3D). This cavity is closed in the closed conformation (Fig. 3E). Importantly, several studies have reported that TMEM16 can induce membrane distortion, reducing membrane thickness in an area near the subunit cavity and thereby lowering the energetic cost for phospholipid movement from one leaflet to the other (Bethel and Grabe, 2016; Falzone et al., 2022; Kalienkova et al., 2019). By calculating the local membrane thickness, we saw that both conformations of the protein induced some membrane thinning in the immediate proximity (within 1 nm) of the protein, and the thinning was more pronounced in the case of the open conformation (Fig. 3F). For the latter, two thinner regions were observed around each protomer, one in the proximity of each subunit cavity, between TM4 and TM6, and the other at the interface between the protomers. In summary, our MD simulations support the scramblase activity of Ist2, and show an inter-relationship between Ist2 activity and the physico-chemical properties of the bilayer.

### Influence of the scramblase domain of Ist2 on cellular lipid homeostasis

Having established that Ist2 can scramble phospholipids, we next investigated whether this activity affects lipid homeostasis in yeast. The cytosolic C-ter tail of Ist2 mediates the transport of PS from the ER to the PM via its interaction with the PS-transfer proteins Osh6 and its homolog Osh7 (D’Ambrosio et al., 2020; Wong et al., 2021). Mutations in the C-ter tail of Ist2 that prevent interaction with Osh6/7 mimic the phenotypes associated with the deletion of Osh6 and Osh7. To assess the defects associated with the deletion of the N-ter TMD of Ist2 independently of the function of its cytosolic tail, we constructed chromosomal yeast mutants expressing truncated versions of Ist2. We used GFP-tagged variants to verify their intracellular localization: a mutant lacking the majority of the TMD except for the last two TM helices, GFP-ΔNIst2, retains the localization at ER-PM contact sites (Manford et al., 2012), whereas Ist2-ΔC lacking the C-ter cytosolic tail localizes throughout the ER (Fig. 4A).

**Figure 4.**
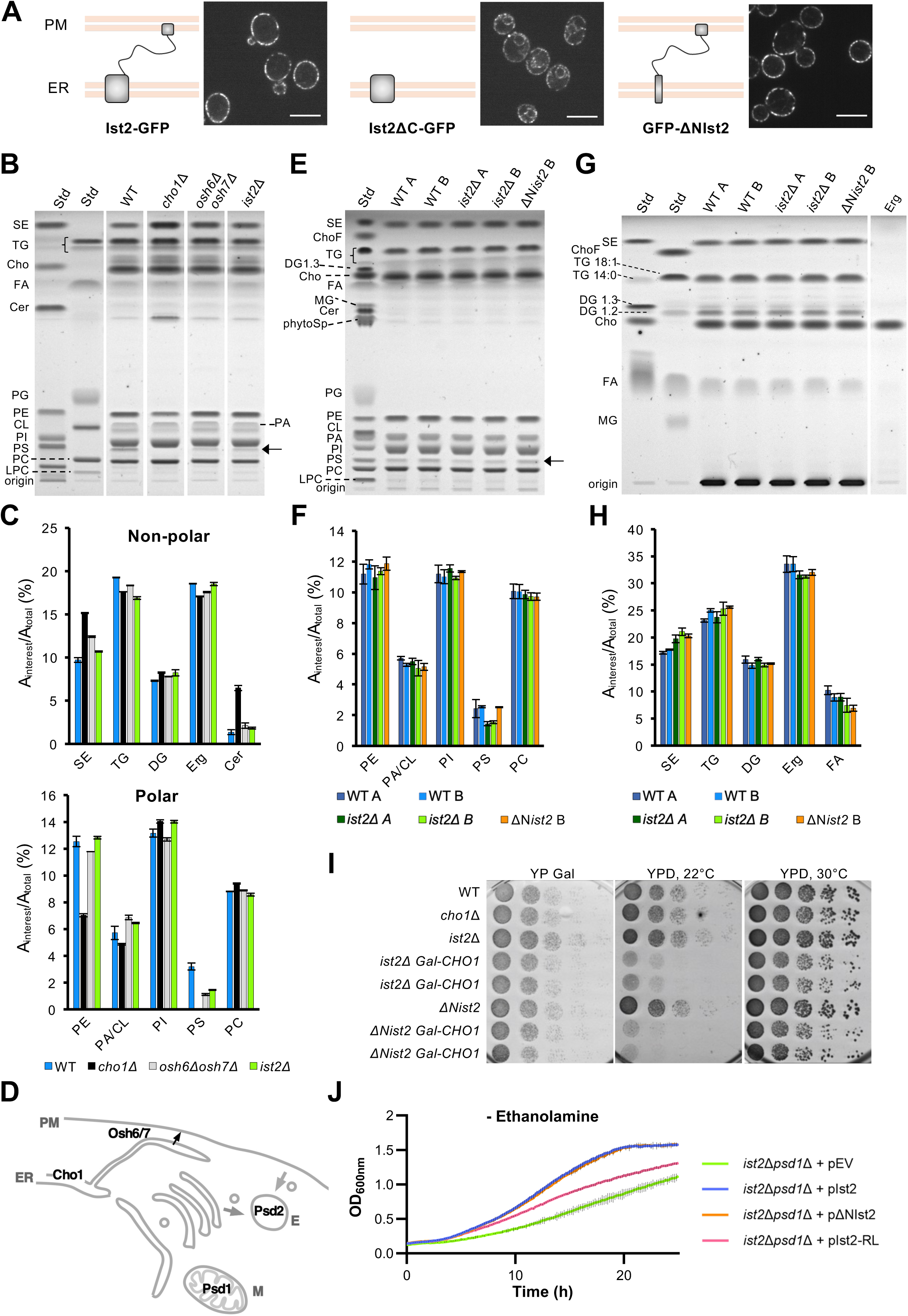
The influence of mutations in Ist2-Nter domain on the cellular lipidome and PS synthesis/decarboxylation. **(A)** Localization of WT and truncated versions of Ist2 tagged with GFP in the chromosomal *IST2* locus. The schematic illustrates Ist2 anchored to the endoplasmic reticulum (ER) via its N-terminal domain and tethered to the plasma membrane (PM) through its C-terminal domain. Corresponding images show Ist2-GFP, Ist2ΔC-GFP and GFP-ΔNIst2 localization. Scale bar = 5 µm. **(B, C)** Thin layer chromatography (TLC) analysis of lipid classes of the indicated yeast strains (B) and lipid classes distribution, separated into neutral lipids (C, non-polar) and phospholipids (C, polar) to facilitate viewing. **(D)** Schematic representation of PS pathways in yeast. PS is synthetized at the ER by the PS synthase Cho1 and can be transported to the PM by the LTPs Osh6/7. PS can be converted to PE by the PS decarboxylase Psd1 at the mitochondria or Psd2 in the endosome. **(E, F)** TLC analysis of lipid classes of the indicated yeast strains (E) and phospholipid class distribution (G). **(G,H)** TLC of neutral lipid classes in yeast strains (G) and lipid class distribution (H). Equal phospholipid masses were applied for each sample in B, E and G. Strains labelled A, or not specified, were grown with agitation and strains labelled B were grown without it. The abbreviations used for standards (Std) refer to: SE: cholesterol ester; TG: triglyceride (18:1 or 14:0); Cho: cholesterol; Erg: ergosterol; DG: diglycerides (1:3 or 1:2); FA: free fatty acid; MG: monoglyceride; Cer: ceramide; Sphin: phytosphingosine; PG: phosphatidylglycerol; PE: phosphatidylethanolamine; CL: heart cardiolipin; PA: phosphatidic acid; PS: phosphatidylserine; PC: phosphatidylcholine; LPC: lysophosphatidylcholine. Graphs show the mean ± range of two independent replicates. **(I)** Growth kinetics of *ist2Δpsd1Δ* cells expressing WT and mutated versions of Ist2, tagged with BFP at N-ter, grown in SC–His medium at 30°C without ethanolamine supplementation. Growth of these strains upon addition of ethanolamine is shown in Fig. S2D. The growth curves represent mean ± range of absorbance at 600 nm (OD_600_) of two independent biological replicates over time (minutes). Each experiment is representative of at least three independent biological replicates. **(J)** Genetic interaction between *IST2* and *CHO1*. The expression of *CHO1* is driven by the Gal promotor. Ten-fold serial dilutions were spotted on galactose plates to allow the expression of *CHO1* and glucose plates to shut down the expression of *CHO1*. Plates were incubated for 3 days at 22°C or 30°C.

We previously showed that despite their impairment in PS transfer from the ER to the PM, *ist2*Δ or *osh6*Δ *osh7*Δ mutants do not show a strong defect in the steady-state cellular distribution of PS, evaluated with the PS probe C2_Lact_–GFP (D’Ambrosio et al., 2020). We did not observe any defect in C2_Lact_–GFP distribution in Δ*Nist2* mutant cells, and a slight defect in *ist2*Δ*C* truncation mutants (Fig. S2A,B). We then used thin-layer chromatography (TLC) to analyze the impact of Ist2 truncations on the total yeast lipidome. As previously shown, the deletion of *IST2* results in a decrease in total cellular PS levels by about 50%, similar to the defect observed in *osh6*Δ *osh7*Δ or in strains with mutations in the Ist2-Cter cytosolic tail (Fig. 4B,C) (D’Ambrosio et al., 2020). This reduction is likely due to the feedback inhibition of the sole yeast PS synthase, Cho1, caused by impaired PS export from the ER. As expected, no PS can be detected in *cho1*Δ cells. Furthermore, *cho1*Δ cells display a two-fold decrease in phosphatidylethanolamine (PE), which can be generated from PS by two PS decarboxylases, Psd1 and Psd2 (Fig. 4B-D) (Lenoir et al., 2021; Storey et al., 2001). Some other lipids are also highly affected in *cho1*Δ cells, especially ceramide (∼4-fold increase) and sterol ester (SE; 30% increase), whereas this is not the case in *osh6*Δ *osh7*Δ or *ist2*Δ strains (Fig. 4B,C). In contrast to *cho1*Δ, these strains also show normal PE levels, as previously reported (D’Ambrosio et al., 2020).

We then compared the lipidomes of *ist2*Δ and Δ*Nist2* cells. In contrast to *ist2*Δ, no reduction in cellular PS levels is observed in Δ*Nist2*. The total levels of other major phospholipids are also not affected (Fig. 4E,F). A separate analysis of more hydrophobic lipids revealed a small but consistent increase (∼15%) in SE in both *ist2*Δ and Δ*Nist2,* as well as a slight decrease in ergosterol levels (Fig. 4G,H). This observation led us to evaluate the distribution of sterols in these strains using the fluorescent sterol probe GFP-D4H (Encinar Del Dedo et al., 2021). A polarized PM-localized GFP-D4H signal was observed in WT as well as in *ist2* mutant strains, in contrast to two control strains, *cho1*Δ and *sac1*Δ, where the GFP-D4H signal was highly perturbed (Fig. S2C). Overall, these results show that deletion of the N-ter scramblase domain of Ist2 does not lead to any gross perturbations of the cellular lipidome.

A more direct way to check for defects in PS export from the ER is using an *in vivo* PS transport assay (D’Ambrosio et al., 2019; Filseck et al., 2015; Maeda et al., 2013). This assay requires the use of *cho1*Δ cells, in which PS production can be restored via acylation at the ER level of exogenously added lyso-PS. However, we were not able to evaluate whether Δ*Nist2* affects PS export because we could not construct a Δ*Nist2 cho1*Δ strain, likely due to a negative genetic interaction between *CHO1* and *IST2.* This interaction is difficult to verify due to the growth and mating defects of *cho1*Δ cells, but could be observed when *CHO1* expression is under the control of a galactose (Gal)-inducible promotor: growth of *ist2*Δ *Gal-CHO1* and of Δ*Nist2 Gal-CHO1* cells is repressed on glucose-containing plates (YPD), whereas all strains grow well when Gal is used as the carbon source (Fig. 4I). Of note, Δ*Nist2* shows a stronger interaction with *cho1* than full deletion of *IST2,* and the effect shows a strong temperature-dependence, being more pronounced at lower temperature (22°C).

Another assay to evaluate the function of Ist2 in Osh6-mediated PS transport is by genetic interaction with *psd1*Δ (Wong et al., 2021). The growth of *ist2*Δ *psd1*Δ cells is strongly inhibited in synthetic media lacking ethanolamine whereby direct synthesis of PE via the Kennedy pathway is blocked (Storey et al., 2001), because supply of PS to the other PS decarboxylase, Psd2, is also diminished (Fig. 4D,I and Fig. S2D). Growth in this strain is rescued by the expression of WT Ist2, but not by the expression of an Ist2 mutant with randomized C-ter tail (Ist2-RL), which cannot interact with Osh6 (D’Ambrosio et al., 2020; Kralt et al., 2014) (Fig. 4I). In contrast, expression of ΔN-Ist2 rescues growth as efficiently as the WT plasmid, suggesting that the TMD of Ist2 is not implicated in this pathway. Finally, we considered the possibility that Ist2 could function as an anion channel, because this activity is shared by a number of other TMEM16 proteins instead or in addition to their scramblase activity (Jan and Jan, 2024; Malvezzi et al., 2013). It has been suggested that the increased sodium tolerance (ist)-phenotype of *ist2*Δ cells (Entian et al., 1999) is linked to a Ca^2+^-activated Cl^-^ channel activity (Kunzelmann et al., 2016). However, this phenotype, revealed on YPD plates supplied with 1 M NaCl, is weak and likewise not dependent on the TMD of Ist2, as it can be observed in *ist2*Δ but not in *ist2*Δ*C* cells (Fig. S2E). In summary, our results so far do not reveal a direct effect of Ist2 TMD on the cellular lipidome and suggest that the scramblase activity of Ist2 is not required to promote efficient PS export via Osh6.

### Genes required for vesicular budding from the ER display negative interaction with Ist2 scrambling mutants

Defects in lipid transport pathways are often difficult to detect (Peter et al., 2022; Toulmay et al., 2022). We decided to evaluate the effect of Ist2 TMD on ER processes that would be directly impacted by trans-bilayer lipid distribution in the ER or by localized changes in the bilayer composition. One such process is the formation of COPII vesicles, which occurs at discrete ER-exit sites where the COPII coat assembles following the activation of the small GTPase Sar1 by the ER-resident GTP-exchange factor Sec12 (Fig. 5A) (Hutchings et al., 2021). Vesicle formation requires deformation of the ER bilayer. It was shown that lyso-phospholipids, which have a conical shape, can promote vesicle formation by inducing membrane curvature, and membrane protein asymmetry can oppose it (Copic et al., 2012; Derganc et al., 2013; Melero et al., 2018).

**Figure 5.**
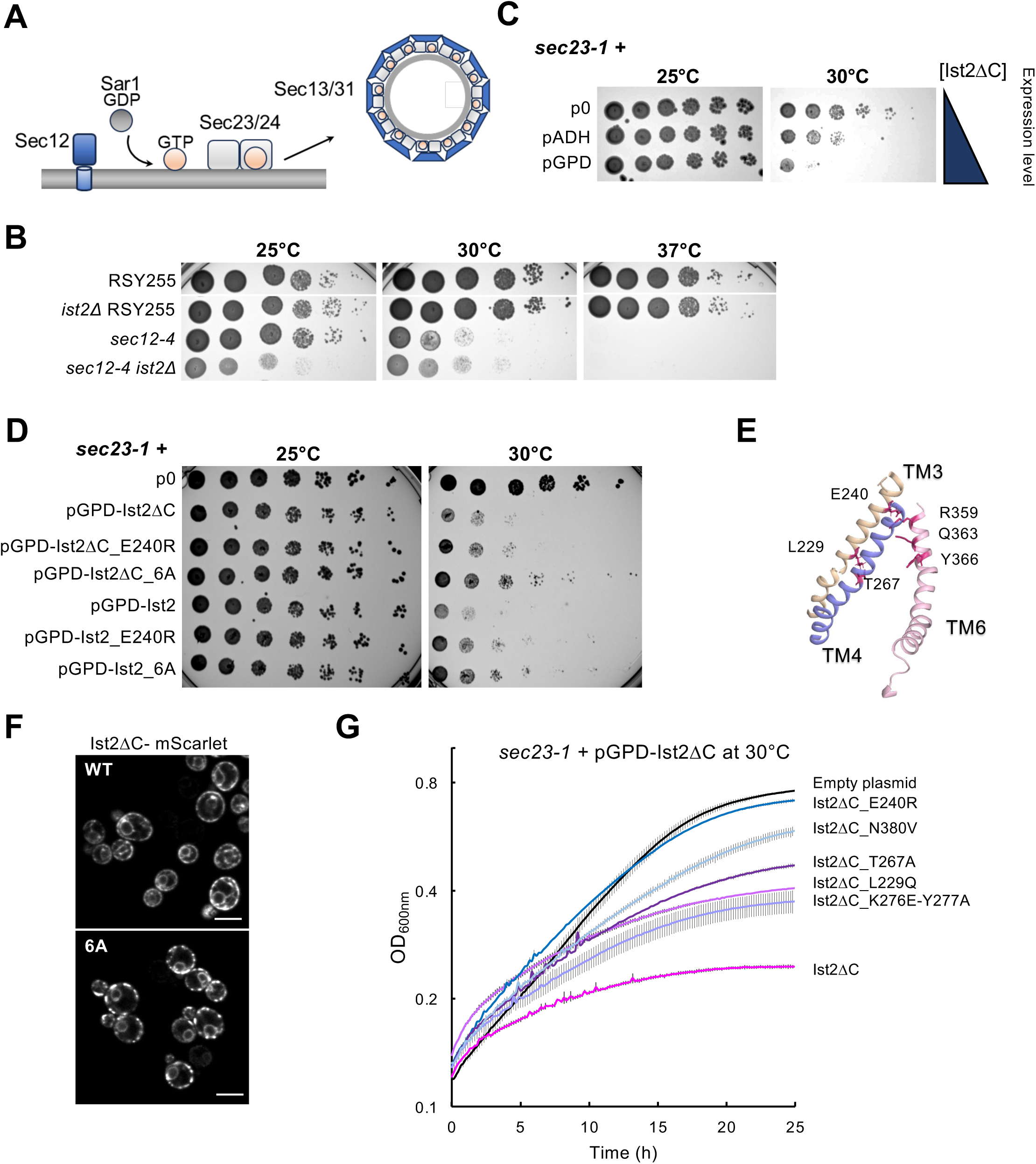
Mutation or overexpression of *IST2* mutants affects the growth of temperature-sensitive (ts) *sec* mutants required for COPII vesicle formation at the ER. **(A)** Schematic simplified of the COPII vesicles pathway. **(B)** Viability assay of the indicated yeast strains grown on YPD agar. Ten-fold serial dilutions of overnight cultures were spotted onto the plates and incubated for 6 days at 25°C, 30°C, and 37°C. **(C)** Viability assay of *sec23-1* cells expressing Ist2ΔC at different levels. Ten-fold serial dilutions of overnight cultures were spotted on SC–URA agar medium and incubated for 6 days at 25°C and 30°C. The triangle represents increase in the levels of Ist2ΔC expression in the *sec23-1*, using an empty plasmid (p0) or plasmids with ADH promoter for intermediate expression and GPD promoter for high expression. **(D)** Viability assay of *sec23-1* cells overexpressing WT, truncated and mutant versions of Ist2 from a centromeric plasmid with a GPD promotor. Ten-fold serial dilutions of overnight cultures were spotted on SC– URA agar medium and incubated for 6 days at 25°C and 30°C. p0 refers to an empty plasmid (pRS316). **(E)** Close-up view of the transmembrane helices TM3, TM4, and TM6 of Ist2 that are implicated in lipid scrambling. Residues L229, E240, T267, R359, Q363, and Y366, conserved with residues involved in lipid scrambling in TMEM16, are displayed as red sticks. These residues were mutated to alanine in Ist2_6A mutants. The rest of the structure is not shown for clarity. **(F)** Localization of Ist2ΔC-mScarlet and Ist2ΔC_6A-mScarlet in *sec23-1* cells. **(G)** Growth kinetics of *sec23-1* cells overexpressing WT and mutated versions of Ist2ΔC, tagged with HaloTag, grown in SC–URA medium at 30°C. The growth curves represent the mean ± range of absorbance at 600 nm (OD_600_) of two independent biological replicates, measured in technical duplicates over time (minutes). Each experiment is representative of at least three independent biological replicates.

Furthermore, COPII binding to membranes *in vitro* is affected by phospholipid composition (Matsuoka et al., 1998). The activity of an ER scramblase could therefore negatively impact the formation of COPII vesicle by reducing the amount of asymmetry between the two leaflets, or it could have a positive or negative feedback by changing the phospholipid composition of the cytosolic leaflet.

To test these possibilities, we assayed for genetic interactions between *ist2* and COPII temperature sensitive (*ts*) mutants (Schekman and Novick, 2004). We could observe a genetic interaction between *sec12-4* and *ist2*Δ. Interestingly, this interaction was temperature-dependent, negative at 25°C and slightly positive at 30°C (Fig. 5B and Fig. S3A). In contrast, *ist2*Δ had no effect on the growth of *sec17-1*, which acts downstream of COPII, blocking vesicle fusion (Fig. S3B). We used another COPII *ts* allele, *sec23-1,* but we had difficulties in combining this mutation with *ist2*Δ. We therefore decided to test the opposite phenotype, i.e., the effect of over-expression of *IST2.* Over-expression of Ist2 TMD (Ist2ΔC) negatively affects the growth of *sec23-1* cells at 30°C in a dose-dependent manner (Fig. 5C), as does the expression of full-length Ist2 (Fig. 5D).

We used the over-expression phenotype in the *sec23-1* background to compare the effect of mutations in Ist2 TMD that could impair its scrambling activity. Six residues of interest in TM3 (L229, E240), TM4 (T267), and TM6 (R359, Q363, Y366) were selected based on their conservation in afTMEM16 and nhTMEM16, where they were shown to be important for phospholipid scrambling (Fig. 1B and Fig. 5E) (Lee et al., 2018). Among these residues, L229, T267 and Y366 were also identified by our MD simulations to be in contact with the lipid headgroup during flip-flop (Fig. 3D). Over-expression of single charge-reversion Ist2 mutants E240R, either as a full-length or C-ter-truncated version, partially improves growth of *sec23-1* cells at 30°C, compared to WT plasmids. The effect is even stronger when all six of these residues are mutated to alanine in the Ist2ΔC_6A and Ist2_6A mutants (Fig. 5D). Addition of a C-ter tag does not impact on the toxicity of Ist2ΔC in *sec23-1* (Fig. S3C). Using a plasmid expressing mScarlet-tagged Ist2ΔC, we verified that mutating the six residues to alanine does not affect protein levels or localization (Fig. S3D and Fig. 5F). We then used a kinetic growth assay to more precisely compare the effect of single or double Ist2ΔC point mutants on growth of *sec23-1* at 30°C. Under these conditions, we observed the strongest effect for E240R, followed by N380V, T267A, L229Q and the K276E_Y277A double mutation (Fig. 5G; K267 and Y277 represent two additional residues identified by simulations). These mutants were expressed at similar levels (Fig. S3E). However, we could not observe any effect of the mutation E240R or N380V on Ist2 scrambling activity *in vitro* (Fig. S3F,G). Nevertheless, these results suggest that the scrambling activity of Ist2-TMD has an impact on the COPII transport pathway at the ER.

### Ist2 scrambling activity affects LDs and other lipid pathways at the ER

Another process that requires ER membrane asymmetry is the formation of LDs that bud from the ER (Chorlay et al., 2019; Huang et al., 2021; Nieto et al., 2023). In agreement, it has been shown in mammalian cells that downregulation or loss of the ER scramblases TMEM41B and VMP1 results in accumulation of cytosolic LDs (Huang et al., 2021; Li et al., 2021; Moretti et al., 2018). Our whole-cell lipid analyses suggested an increase in neutral lipids in *ist2*Δ and in Δ*Nist2* mutants (Fig. 4). In agreement, we saw a small but significant increase in the number of LDs present in *ist2*Δ (+26%),Δ*Nist2* (+21%) and in *ist2_6A* mutant cells (+14%), but not in *ist2*Δ*C* (Fig. 6A,B). Furthermore, given that phospholipid synthesis is restricted to a single ER leaflet, we reasoned that a decrease in ER scrambling activity could affect the phospholipid composition of the LD monolayer. To probe for changes in LD surface composition, we decided to use an amphipathic helix (AH) derived from the human LD protein perilipin 4 (PLIN4), which targets LDs in yeast. Previously, we generated a series of PLIN4 AH mutants with subtle changes in their amino acid composition that impacted LD targeting (Copic et al., 2018). One of these AH variants, [2Q, 2V], composed of 4 x 33 aa stretches of the same sequence, interacts weakly but specifically with LDs in yeast and is overall negatively-charged (Fig. 6C) (Copic et al., 2018). We reasoned that the weak LD affinity and negative charge of PLIN4-AH[2Q,2V] should make it sensitive to changes in LD surface composition, in particular to changes in electrostatics resulting from an increase or decrease in charged lipids such as PS. In WT cells, this probe localized to LDs in a fraction of cells. Interestingly, its localization to LDs increased in *ist2*Δ, Δ*Nist2* and *ist2_6A* mutant cells, whereas it decreased in *ist2*Δ*C,* and to a lesser extent in *osh6*Δ*osh7*Δ (Fig. 6D,E), possibly reflecting a decrease or an increase in surface charge, respectively. Overall, these results suggest that Ist2 influences the formation and composition of LDs.

**Figure 6.**
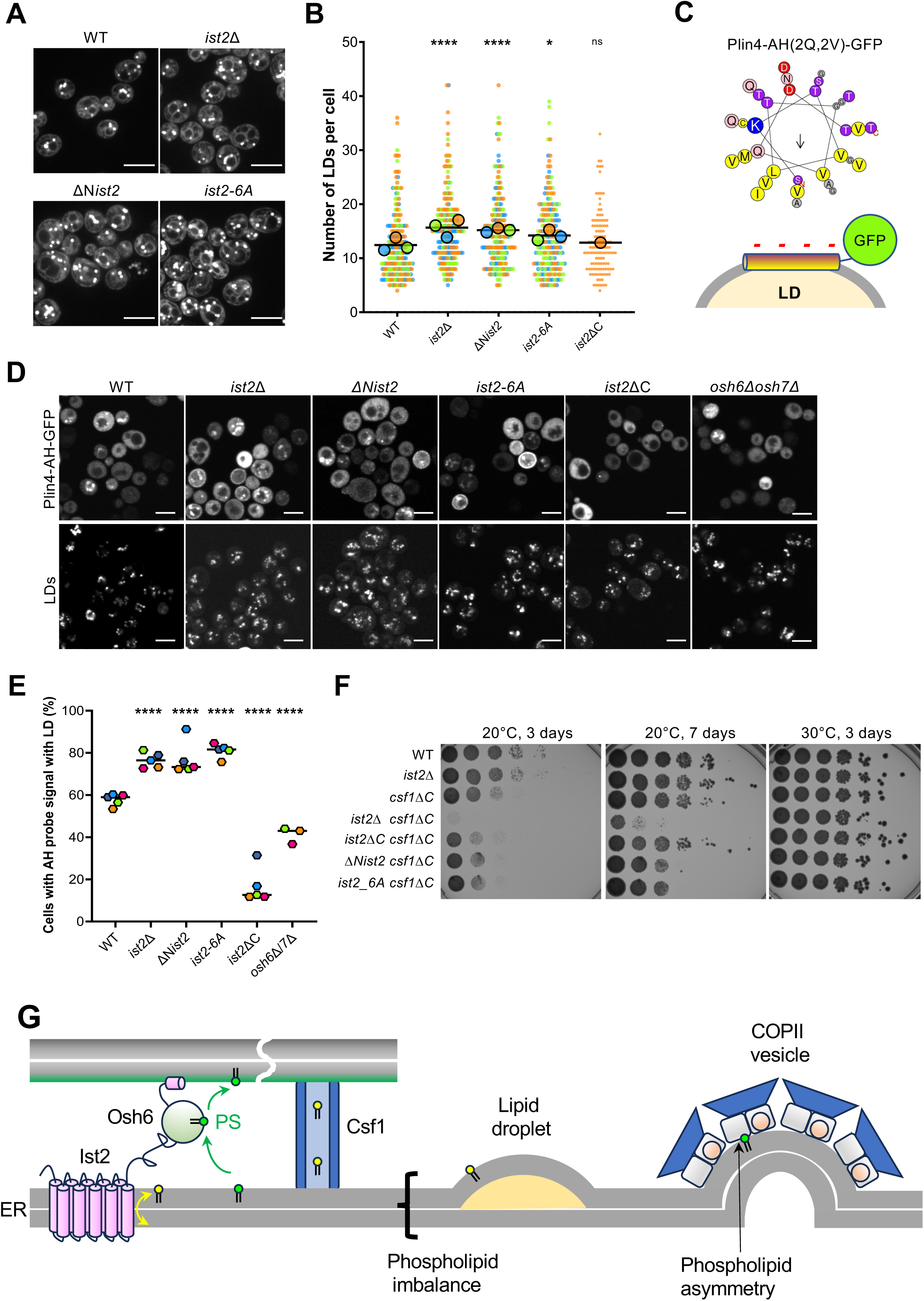
The influence of *ist2* scrambling mutants on LDs and lipid transport. **(A)** Distribution and abundance of LDs stained with Bodipy in WT cells and cells with chromosomal mutations in *IST2*, as indicated. Scale bar = 5 µm. **(B)** Quantification of the number of LDs per cell in indicated strains (n≥60, N=3). The significance was assessed by Kruskal-Wallis and Dunn’s post-hoc test, P>0.05 (ns), P<0.01 (**), P<0.0001 (****). **(C)** Illustration of the PLIN4-AH probe [2Q,2V], plotted using Heliquest (Gautier et al., 2008). The probe contains 4 repeats of the indicated sequence of 33 amino acids, fused to GFP (Copic et al., 2018). **(D)** Distribution of the AH probe in indicated yeast strains bearing chromosomal mutations in *IST2* (left panels), with total LD staining using AutoDot shown in right panels. Scale bar = 5 µm. **(E)** Quantification of percentage of cells with the AH probe localized to LDs. Data are representative of at least two independent biological replicates (n≥100). The significance was assessed by Kruskal-Wallis and Dunn’s post-hoc test, P>0.05 (ns), P<0.01 (**), P<0.0001 (****). **(F)** Viability assay of the indicated yeast strains. Ten-fold serial dilutions of overnight cultures were spotted on YPD agar medium and incubated for 3 and 6 days at 20°C and 30°C. **(G)** Schematic representation of processes that are affected by phospholipid transport activity mediated by Ist2, either scrambling of phospholipids (yellow) or transfer of PS (green).

Finally, we explored whether Ist2 activity might influence other lipid transport pathways. Scramblases have been shown to play an important role in the formation of autophagosomes, where their activity is coupled to lipid transfer by channel-like LTP of the Vps13 family, such as Atg2. However, we did not detect any defect in general autophagy in *ist2*Δ cells, in agreement with a recent report (Fig. S3H,I) (Liu et al., 2024). We instead focused on *CSF1*, which encodes another Vps13-like LTP that can channel phospholipids between the ER and other compartments (Peter et al., 2022; Toulmay et al., 2022). As suggested by a genome-wide screen (Costanzo et al., 2016), we could confirm a cold-temperature dependent negative genetic interaction between *ist2*Δ and a null allele of *CSF1, csf1*Δ*C* (Peter et al., 2022) (Fig. 6F). In contrast, *ist2*Δ*C* had no effect on the growth of *csf1*Δ*C*, whereas the scramblase mutants Δ*Nist2* and the *ist2-6A* had some effect, but less than *ist2*Δ, again showing non-additive effects between deletions of the two subdomains of Ist2. Together, our data suggest that Ist2-TMD impacts several processes originating from the ER, and these effects correlate with mutations in residues implicated in phospholipid scrambling (Fig. 6G). We therefore propose that phospholipid scrambling by Ist2 affects lipid pathways at the ER.

## Discussion

We show that the budding yeast protein Ist2 possesses a non-selective phospholipid scrambling activity in its ER-localized TMD (Fig. 6G). This is not surprising, given the conservation between this domain and the TMEM16 protein family, in particular its closest fungal orthologs nhTMEM16 and afTMEM16, which have both been demonstrated to act as phospholipid scramblases (Brunner et al., 2014; Falzone et al., 2019). Ist2 is the only TMEM16-like protein present in budding yeast, and no dedicated ER scramblase has been identified to date in yeast. Our evidence is based on the reconstitution of Ist2 that was expressed and purified from its native host using mild detergent to preserve its function and the integrity of reconstituted proteoliposomes. We were able to detect robust lipid scrambling both with full-length Ist2 and with Ist2ΔC, lacking the C-ter cytosolic tail, which is unique to yeasts. Importantly, similar to TMEM16 and other scramblases evaluated so far (Sebinelli et al., 2024), Ist2 can scramble a variety of different phospholipid substrates and does not show a preference for a specific lipid headgroup. These results were confirmed by MD simulations, where we used both a simple and a complex, ER-like bilayer composition (Reinhard et al., 2024). Interesting, Ist2 activity *in silico* was affected by bilayer composition and showed preference for PC, PS and PE over PI and PA, as well as for shorter acyl chains).

Both *in vitro* and *in silico*, phospholipid scrambling by Ist2 was not sensitive to Ca^2+^. This deviates from the activity of other TMEM16 scramblases, which are all activated by Ca^2+^ (Bushell et al., 2019; Falzone et al., 2019; Kalienkova et al., 2019; Malvezzi et al., 2013), but is in accordance with the lack of conservation of Ca^2+^-binding residues in the Ist2 sequence. It is possible that Ist2 is a constitutive ER scramblase, or it could be regulated by a different mechanism. Our AF2 and trROSETTA-predicted structural models suggested the possible occurrence of both closed and open conformations for the Ist2 subunit cavity in the absence of Ca^2+^, pointing to alternative mechanisms to Ca^2+^ binding for the control of phospholipid scrambling. These questions will be the subject of further investigations.

The TMEM16 scramblases are suggested to function via a ‘credit card’ mechanism (Pomorski and Menon, 2006), whereby the phospholipid headgroups sliding through the lipid bilayer utilize a hydrophilic slit between TM helices 3-7 that form the subunit cavity. This mechanism was observed in our MD simulations, and mutagenesis of some of the residues lining this pathway resulted in a reversal of the phenotypes in yeast. Alternative or additional mechanisms of lipid scrambling have also been proposed (Jan and Jan, 2024). Both afTMEM16 and nhTMEM16 can induce membrane thinning (Bethel and Grabe, 2016; Falzone et al., 2022; Kalienkova et al., 2019), which would facilitate the passage of phospholipids through the membrane; we have likewise observed membrane thinning in our MD simulations, particularly with the ‘open’ (active) model of Ist2. Furthermore, ‘out-of-the-groove’ scrambling has also been observed for both of these proteins, whereby phospholipids utilize an alternative pathway that does not require Ca^2+^ activation (Falzone et al., 2022; Feng et al., 2024). Out-of-the-groove scrambling of nhTMEM16 is highly dependent on membrane lipid composition (Feng et al., 2024). Unlike Ist2, afTMEM16 and nhTMEM16 both function at the PM, therefore in a very different lipid environment. These considerations are all important when evaluating the activity of Ist2 in comparison to these highly-studied scramblases, and can guide further design of Ist2-scrambling mutants.

In cells, direct evidence of scramblase activity is difficult to obtain. The limitation of bulk cellular lipid assays performed at steady state is that they cannot detect localized effects and that a defect in one pathway can be compensated by other pathways or by other lipids (Bao et al., 2021; Peter et al., 2022). Furthermore, while no other dedicated ER scramblase has so far been described in yeast, many proteins have been suggested to contribute to the scramblase activity of the ER, such as GPCRs (Goren et al., 2014) or all ER protein insertases (Li et al., 2024). Nevertheless, the phenotypes that we could observe upon deletion or over-expression of the Ist2 TMD strongly support a scramblase activity for this domain (Fig. 6G). Our strongest phenotype is the effect of Ist2 TMD over-expression on the COPII vesicular pathway. The propensity of a cellular membrane to be deformed by a coat machinery depends on the efficient recruitment of the coat components and on physical factors including membrane bending modulus, membrane tension, and protein crowding. In the case of the COPII coat, lipid unsaturation and negative charge have been shown to facilitate the recruitment of the small G protein Sar1 and of the Sec23/24 complex, respectively (Matsuoka et al., 1998), whereas cargo crowding on the luminal side has been shown to oppose the mechanical effect of the coat (Copic et al., 2012; Derganc et al., 2013). However, the difficulty in making liposomes with asymmetric lipid composition has hampered testing the role of lipid distribution between the two leaflets on coat-induced membrane budding. Nevertheless, indirect evidence suggests an impact of interleaflet lipid distribution on membrane deformation. In Drosophila, high scramblase activity leads to a symmetric and highly-deformable PM (Shiomi et al., 2021). Furthermore, a recent study in mammalian cells has shown that TMEM41B influence export of GPI-anchored protein from the ER (Cao et al., 2023). Finally, molecular dynamics studies suggest that polyunsaturated lipids, when present in the convex side, facilitate membrane deformation (Pinot et al., 2014; Tiberti et al., 2020). By contrast, recent biophysical measurements indicate that asymmetric membranes containing PE and PS in one leaflet are rather stiff (Frewein et al., 2023). Clearly, the link between membrane asymmetry and coat-induced membrane deformation is a challenging issue for future studies.

More well-documented is the influence of mammalian ER scramblases on the production of LDs or lipoprotein particles that bud into the ER lumen, which appears particularly pronounced in strongly-secretory liver cells or tissue (Huang et al., 2021; Wu et al., 2024). In agreement with these observations, deletion of Ist2 TMD or substitution of its conserved scrambling residues both affect LD abundance and composition to a similar extent. Another important clue is that a number of our cellular assays showed a strong dependence on temperature, suggestive of changes in membrane composition upon Ist2 TMD deletion and a homeoviscous membrane adaptation (Klose et al., 2012; Renne and Kroon, 2017). Together, these observations support the implication of Ist2 TMD domain in the regulation of the ER lipid homeostasis *via* its phospholipid scramblase activity, in agreement with our *in vitro* and *in silico* data. A scramblase activity of Ist2 TMD can also explain other phenotypes that have been reported for *ist2*Δ cells, such as osmotic sensitivity (Smith et al., 2024) and an effect on ER-phagy (Liu et al., 2024).

An unusual feature of Ist2 is the presence of a long cytosolic tail that interacts with the PM, localizing Ist2 to ER-PM contact sites (Fig. 6G). Although this tail is unique to yeasts, two other mammalian TMEM16 proteins, TMEM16H and TMEM16K, have been shown to localize to the ER and engage in contacts with the PM and the endosomes, respectively (Jha et al., 2019; Petkovic et al., 2020). TMEM16K has also been shown to act as a scramblase. These observations raise the obvious question of the advantage of localizing an ER lipid scramblase to a membrane contact site. We showed previously that the Ist2 tail interacts with the PS-specific LTP Osh6 and is required for efficient transport of PS from the ER to the PM (D’Ambrosio et al., 2020). Mutations in the tail or deletion of Osh6 and its homolog Osh7 result in a decrease in the total levels of PS, which can be explained by a feed-back inhibition of the PS synthase Cho1 at the ER (Kannan et al., 2017; Sohn et al., 2016). Cho1 has generally been assumed to synthesize PS in the cytosolic leaflet of the ER (Chauhan et al., 2016), although this has been recently put under question by the structural characterization of the human PS synthase PSS1, whose active site is positioned in the ER lumen (Long et al., 2024). Under either scenario, the scramblase activity would affect the total pool of PS available for transfer by Osh6, and thereby would be expected to modulate the efficiency of PS export from the ER. The intrinsic set-up of Ist2 would allow for direct coupling between two lipid transport activities, lipid flip-flop and lipid transfer between compartments, enabling more precise regulation of the two processes and a faster and non-linear response (feed-back mechanisms). However, we have not detected any impact of the deletion of the TMD of Ist2 on the cellular PS levels or distribution, in contrast to the deletion of the C-ter tail. One issue may be the sensitivity of our assays, especially since we could not perform the cellular PS transport assays due to the synthetic interaction between *cho1*Δ and Δ*Nist2*.

Despite the lack of direct evidence so far, our results reveal some important clues suggesting that the two domains of Ist2 function in a coordinated manner. A number of our assays revealed non-additive effects between the ΔN and ΔC deletions of Ist2. This was evident in the genetic interaction with cho1Δ, where ΔNist2 had the strongest impact, or in the interaction with csf1Δ, where the interaction with *ist2*Δ was stronger than the sum of the ΔN and ΔC interactions, as well as in the LD phenotypes, where the effect of ist2ΔC contrasted the effects of *ist2*Δ and Δ*Nist2.* One exciting possibility is functional interplay between the TMD and the C-terminal tail in the Ist2 molecule, which we are now in position to explore *in vitro* by combining the lipid scrambling and lipid transfer function assays.

## Acknowledgments

We wish to acknowledge Sophie Dupré, Anne Claude Gavin, Elizabeth Miller, Simonetta Piatti Elizabeth Venkoff for gift of strains and plasmids, Daniel Lévy and Aurélie Di Cicco for help with reconstitution in proteoliposomes, Juliette Martin for support with the MODELLER software, Cyril Moulin, Ivan Fayol and Maxime Bello for help with strain construction and phenotype testing, Clément Benedetti for assistance with image analysis, and Thibaud Dieudonné, Guillaume Drin, and Bruno Antonny for helpful discussions and comments on the manuscript. We thank the joint IGMM-CRBM “yeast media and technologies service”. This work benefited from the CryoEM platform of I2BC, supported by the French Infrastructure for Integrated Structural Biology (FRISBI) [ANR-10-INSB-05-05], the MRI imaging facility, a member of the national infrastructure France-BioImaging, supported by the French National Research Agency (ANR-10-INBS-04, Investissements d’avenir). All MD simulations were performed at French supercomputing centers (CINES and TGCC) supported by *Grand Equipement National de Calcul Intensif* (GENCI, grant number A0120710138, A0140710138). We acknowledge CC-IN2P3 (https://cc.in2p3.fr) for computing services (data storage and backup). LM acknowledges funding by the *Institut National de la Santé et de la Recherche Médicale* (INSERM). This work was supported by the Agence Nationale de la Recherche (ANR-20-CE13-0030-02, ANR-21-CE11-0015-01 and ANR-23-CE44-0026).

## Materials and Methods

### Plasmid construction

All plasmids used in this study are listed in Table S1. The Gibson Assembly Hifi Kit (Ozyme) was used to generate the following plasmids: pADH-Ist2(N), pGPD-Ist2(N), pGPD-Ist2, and their corresponding tagged versions (mScarlet or Halo tag amplified from Addgene 179067 and 188929 plasmids, respectively) (Bean et al., 2022). Site-directed mutagenesis kit (Agilent) was used for generating point mutations in plasmids. Briefly, plasmids were amplified using overlapping oligonucleotides containing the desired mutations, designed with the most frequent codon usage. All constructs were verified by DNA sequencing. For overexpression of Ist2 in *S. cerevisiae*, the tagged genes were cloned into the pYeDP60 plasmid (Pompon et al., 1996). The preparation of these plasmids was performed as follows. The gene encoding Ist2 was amplified from yeast genomic DNA. The *IST2* gene was inserted into the shuttle pJET1.2 plasmid (Thermo Fisher Scientific) to obtain the pJET-*IST2* plasmid. For the cloning of various PCR fragments into the pYeDP60 vectors, the ‘in-fusion’ reaction (Takara) was used. All the inserted fragments were checked by DNA sequencing.

### Reagents for purification and *in vitro* reconstitution

Products for yeast and bacteria cultures were purchased from Difco (BD Biosciences) and Sigma. *n*-dodecyl-β-d-maltopyranoside (DDM, D310) was purchased from Anatrace. Streptavidin-sepharose resin was purchased from GE/Cytiva (17511301). 1-palmitoyl-2-oleoyl-*sn*-glycero-3- phosphocholine (POPC), 1-palmitoyl-2-oleoyl-*sn*-glycero-3-phosphoserine (POPS), 1-palmitoyl- 2-{12-[(7-nitro-2-1,3-benzoxadiazol-4-yl)amino]dodecanoyl}-sn-glycero-3-phosphoserine (ammonium salt) (NBD-PS), 1-palmitoyl-2-{12-[(7-nitro-2-1,3-benzoxadiazol-4- yl)amino]dodecanoyl}-sn-glycero-3-phosphocholine (NBD-PC), 1-palmitoyl-2-{12-[(7-nitro-2-1,3- benzoxadiazol-4-yl)amino]dodecanoyl}-sn-glycero-3-phosphoethanolamine (NBD-PE) were purchased from Avanti Polar lipids. mCherry was detected using a rabbit anti-mCherry antibody from Invitrogen (PA5-34974). Goat anti-rabbit HRP-coupled IgG antibody (1706515), Bio-Beads® SM-2 Adsorbent (152-3920), and Precision Plus Protein Standards (1610393) were purchased from Biorad. EDTA-free protease inhibitor cocktail (S8830) was purchased from Sigma.

### Expression of Ist2 in yeast membranes

W303.1b/*GAL4* (*a*, *leu2-3*, *his3-11*, *trp1-1:TRP1-GAL10-GAL4*, *ura3-1*, *ade2-1*, *can r*, *cir +*) yeast strain was transformed by the lithium-acetate method. Yeast cultures and recombinant protein expression were performed as previously described (Dieudonné et al., 2022). Yeast cells were then harvested by centrifugation, washed with ice-cold deionized H_2_O, then with ice-cold TEKS buffer (50 mM Tris-HCl pH 7.5, 1 mM EDTA, 0.1 M KCl, 0.6 M sorbitol), and resuspended in TES buffer (50 mM Tris-HCl pH 7.5, 1 mM EDTA, 0.6 M sorbitol) supplemented with protease inhibitors (SIGMAFAST EDTA-free protease inhibitor cocktail) and 1 mM PMSF. The cells were subsequently broken with 0.5 mm glass beads using a ‘Pulverisette 6’ planetary mill (Fritsch). The crude extract was spun down at 1,000 *g* for 20 min at 4 °C to remove cell debris and nuclei. The resulting supernatant was centrifuged at 20,000 *g* for 20 min at 4 °C, yielding S2 supernatant and P2 pellet. The S2 supernatant was then ultracentrifuged at 125,000 *g* for 1 hr at 4 °C. The resulting P2 and P3 pellets were then resuspended at about 30–50 mg.mL^-1^ of total protein in TES buffer.

### Purification of Ist2 from yeast membranes

Membranes obtained after expression of Ist2 (P2 or P3) were diluted to 5 mg.mL^-1^ of total protein in ice-cold buffer A (50 mM MOPS-Tris at pH 7.0, 500 mM NaCl, 20% (w/v) glycerol), supplemented with 1 mM PMSF and protease inhibitors. n-dodecyl-β-D-maltoside (DDM) was then added to 15 mg.mL^-1^, resulting in a DDM/protein ratio of 3/1 (w/w). The suspension was then stirred gently on a wheel for 1 hr at 4 °C. Insoluble material was pelleted by centrifugation at 100,000 *g* for 1 hr at 4 °C. The supernatant was applied onto streptavidin-sepharose resin (1 mL per 3 mg Ist2) and incubated for 2 hours at 6 °C to allow binding of BAD-tagged Ist2mC to the resin. The resin was then washed thrice with three resin volumes of ice-cold buffer A supplemented with 0.5 mg.mL^-1^ DDM. Elution was performed by addition of 60 μg of purified TEV per mL of resin and overnight incubation at 6 °C. Before reconstitution, the eluted fraction was concentrated to 0.3-0.4 mg.mL^-1^ using a Vivaspin unit (100 kDa MWCO).

### Liposome preparation and protein reconstitution

Liposomes were formed from a 9:1 (mol/mol) mixture of POPC:POPS (Avanti Polar Lipids). Lipids were dissolved at 10 mg.mL^-1^ in 5 mL chloroform and dried in rotavapor for approximately 30 minutes under pressure to form a thin lipid film in a glass balloon. After chloroform evaporation, the lipid film was placed under vacuum in a desiccator for at least one hour. Lipids were then suspended in 12.5 mL of buffer R (200 mM NaCl, 50 mM MOPS/Tris, pH 7.0), yielding multilamellar vesicles at a final concentration of 4 mg.mL^-1^. The vesicles were aliquoted, flash frozen in liquid nitrogen and stored at -80°C. Immediately before reconstitution, 400 μL of the vesicles were thawed and allowed to equilibrate at room temperature. All subsequent steps were performed at room temperature. The vesicles were then solubilized at room temperature for 15 minutes with Triton X-100 (TX-100) under agitation using a magnetic stirrer. The detergent:lipid ratio used was 2.5:1 (w/w) as determined by (Rigaud and Lévy, 2003), resulting in final concentrations of 7 mg TX-100.mL^-1^ and 2.7 mg lipids.mL^-1^. Then, 0.3 mol% of C12-NBD-labelled lipids (NBD-PS, NBD-PE or NBD-PC) resuspended in 0.5 mg.mL^-1^ DDM were added to the lipid/TX-100 mixture, along with 1 mM EGTA. The purified protein was added to the NBD-labelled detergent/lipid suspension at 14.4 mg.mmol^-1^ protein/lipid ratio (∼ 0.05 mg.mL^-1^ protein) and incubated for 30 min at room temperature. Triton X-100 and DDM were removed by adding pre-washed Bio-beads SM-2 adsorbent (Bio-Rad) in three steps: first, the sample was incubated for 2 hours with 10 mg Bio-beads/mg TX-100, then a second aliquot of Bio-beads was added (10 mg/mg TX-100) and the sample was incubated for another 2 hours. Lastly, Bio-beads from the first two steps were removed and an additional 20 mg Bio-beads/mg was added for 1 hour to the sample (Rigaud and Lévy, 2003). Finally, the proteoliposomes were separated from the beads and stored at 4°C for up to one week.

### Sucrose gradient fractionation

Reconstitution was assessed using a discontinuous sucrose gradient, prepared with five sucrose solutions (5, 10, 20, 30 and 40%) prepared in buffer R. Each solution was layered in 2 mL increments in a centrifuge tube. Then, 350 µL of samples were carefully placed on top of the 5% sucrose layer and spun down at 160,000 g for 20 hours. After centrifugation, 500 µl fractions were collected from the top to evaluate lipid, protein and sucrose concentration. Sucrose concentration was measured with a hand refractometer, protein incorporation checked by Western-blotting using an anti-mCherry antibody and lipids were tracked by NBD-fluorescence. For the dithionite transport assay of proteoliposomes separated on a sucrose gradient, 250 µL fractions were collected from the top of the gradient and submitted to dithionite quenching.

### Cryo-EM of proteoliposomes

Samples were prepared with 3 µL of liposomes or proteoliposomes diluted 2-fold in buffer R, spotted onto Lacey grids and vitrified by plunge-freezing performed with an in-house guillotine. Blotting time and force were adjusted on a per sample basis. Images were acquired with a Tecnai Spirit 120 kV LaB6 FEI microscope operated at 120 kV equipped with a K2 Base camera (Gatan).

### Scramblase assay

A 20-µL aliquot of either liposomes or proteoliposomes was diluted to a final volume of 2 mL in buffer R supplemented with either 1 mM EGTA (8 nM free Ca^2+^ concentration) or without EGTA and with 1 mM CaCl_2_. Then, the fluorescence of NBD lipids was monitored over time in a quartz cuvette, using excitation at 470 nm and emission at 530 nm. After 200 seconds, freshly prepared sodium dithionite was introduced at a final concentration of 40 mM and the bleaching of NBD-lipids was followed for ∼800 s. At t=900 s, 0.1% (w/w) TX-100 was added to solubilize the vesicles and to allow complete reduction of the NBD. Data acquisition was done using a Fluorolog®-3 Spectrofluorometer (HORIBA Instruments Incorporated) packed with FluorEssence™ software. All fluorescent points were normalized (*F*_norm_) to the background fluorescence using the following formula: *F*_norm_ = (*F* − *F*_end_)/(*F*_start_ − *F*_end_) ⨉ 100, where *F* is the intensity at each time point, *F*_start_ is the intensity just before dithionite addition and *F*_end_ is the intensity after TX-100 addition.

### Iodide Collisional Quenching

A 10-μL sample of liposomes or Ist2 proteoliposomes was diluted to a final volume of 2 mL in Buffer R supplemented with 40 mM Na_2_S_2_O_3_ (to preserve iodide ions in the solution) and concentrations of KI ranging from 0 to 0.2 M. KCl was used to adjust the ionic strength to 0.2 M. NBD-fluorescence was followed by setting excitation and emission wavelengths at 470 nm and 530 nm, respectively. The mean fluorescence value recorded over the first 5 minutes was taken and used to plot Δ*F* (*F*_0_-*F*). The data were analysed according to the modified Stern-Volmer equation (Vehring et al, 2007; Goren et al, 2014): *F*_0_/Δ*F* = (1/*f_a_* . *K*[*Q*]) + (1/*f_a_*) where *F*_0_ is the fluorescence intensity in the absence of the quencher, Δ*F* is the fluorescence intensity in the presence of the quencher at concentration [*Q*] subtracted from *F*_0_, *f_a_* represents the fraction of fluorescence accessible to iodide ions and *K* is the Stern-Volmer quenching constant.

### Molecular models of the protein

Since there is no crystallography or cryo-EM structure for the yeast Ist2 protein in the protein data bank (PDB), homology modeling techniques were employed to predict 3D models for the transmembrane domain (TMD) of Ist2. HHpred (pairwise comparison of profile hidden Markov models (HMMs)), an online enactment of HHsearch at the Tuebingen toolkit (Gabler et al., 2020), was used for profile-profile searches for TMD of Ist2 homologs in “PDB_mmCIF70” database, using default parameters. HHpred revealed 25%, 24%, 20%, and 16% identities between the TMD amino acid sequence of Ist2 and afTMEM16 (PDB ID: 6E1O), nhTMEM16 (PDB ID: 4WIS), hTMEM16K (PDB ID: 5OC9), and mTMEM16A (PDB ID: 5OYB), respectively.

We used locally installed AlphaFold2 (Jumper et al., 2021) with default structure prediction pipeline, where three different Multiple Sequence Alignments (MSAs) generated by searching the Big Fantastic Database (BFD), MGnify and Uniref90 with jackhmmer from HMMER3.3.2 (Eddy, 2011). During modelling, relaxation was turned off. The locally installed AlphaFold2 gave us homodimer of TMD with open conformation, similar to the crystal structure of nhTMEM16 (PDB ID: 4WIS). In order to sample alternative conformations of TMD, ColabFold with different MSA sizes were used by modifying the "max_msa_clusters" and "max_extra_msa" parameters. We obtained different conformations of TMD, but none of them represented the closed state of TMD. Then we modelled Ist2 TMD using I-TASSER and trRosetta web servers, using fully automated protein structure prediction pipeline. Among all generated models, only models provided by trRosetta tend to be very similar to the closed state conformation. Each model’s similarity to the experimentally open (PDB ID: 4WIS) and closed (PDB ID: 6QMB) conformation were quantified using TM-score. MODELER v10.1 (Webb and Sali, 2016) was then used to build the homodimer model of the closed state using the PDB ID: 6QMB as template. Two trRosetta made promoters were superimposed onto 6QMB structure (RMSD=1.101 Å) using the UCSF Chimera software (Pettersen et al., 2021) (MatchMaker tools) and used as input for the MODELER program. One hundred models were generated, and the one with the lowest discrete optimized protein energy (DOPE) score was adopted for MD simulations. In conclusion, two TMD models were built, representing the “closed” obtained from trRosetta and the “open” state obtained from AlphaFold2. Both models were used in subsequent MD simulations.

### All-atom Molecular Dynamics simulations

Prior to MD simulations, PROPKA 3.2 (Olsson et al., 2011) was used to determine the protonation states of ionizable residues. As a result, standard protonation states at neutral pH were assigned to all residues, except for protonated H506, which are located at the dimer interface and form salt bridges with residues E505 of the other protomer. For both the open and closed structural model, we built a Ca^2+^-bound and a Ca^2+^ unbound form, hence 4 systems were built. Each protein model was embedded into two different lipid mixtures mimicking the ER composition using the membrane builder module of CHARMM-GUI server (Lee et al., 2016). Lipids were symmetrically distributed between the leaflets. For details of bilayer compositions, see Suppl. Table 1. The TIP3P water model was used to solvate the system, and 0.15 M KCl were added to neutralize the system. The resulting systems contained ∼707,000 atoms in total and extended by 21 nm*21 nm*18 nm in XYZ directions.

We carried out MD simulations using the GROMACS 2019.2 molecular dynamics program package with the CHARMM36m all-atom force field. Periodic boundary conditions were applied in all directions. Non-bonded interactions were calculated using the Verlet cutoff scheme, and a buffer tolerance of 0.005 kJ/mol; a cut-off of 12 Å was applied for non-bonded interactions. The particle mesh Ewald (PME) approach was employed for long-range electrostatic interactions, with a Fourier grid spacing of 1.2 Å. The LINCS algorithm was used to constrain bond lengths involving the hydrogen atoms.

After standard energy minimization using the steepest descent algorithm, the resultant systems were pre-equilibrated for 250 ns in isothermal-isobaric ensemble (NPT), in this step backbone atoms of protein were kept fixed with a harmonic positional restraints potential of 2000 kJ mol^−1^ nm^−2^ which allowing other molecules to adjust with protein. Following pre-equilibration, all the systems were equilibrated for 250 ns with a harmonic positional restraints potential of 50 kJ mol^-1^nm^-2^ (in the same NPT ensemble). The temperature was held at 298 K with a v-rescale thermostat (Bussi et al., 2007), using separate temperature coupling for the solvent and solute (protein and membrane). The time constant for temperature coupling was set to 1 ps. During equilibration, the pressure was kept constant using the semi-isotropic Berendsen barostat (Bernetti and Bussi, 2020)] set to 1 bar using a 10 ps coupling constant and compressibility 4.5 × 10^-5^ bar^-1^. The equations of motion were integrated using the leap-frog algorithm, with an integration time step of 2 fs.

Production runs were performed for 1 µs with the same parameters as the equilibration described above, except that the pressure was maintained at 1 bar using the semi-isotropic Parrinello−Rahman barostat (τ_P_ = 10 ps and compressibility = 4.5 × 10^-5^ bar^-1^). In addition, cylindrical (in parallel to the z-axis) flat-bottomed restraints were applied on protein atoms in lipid bilayer, using a force constant of 2000 kJ mol^−1^ nm^−2^, and a cylinder radius of 9.8 nm, to prevent the protein from crossing the periodic boundary and move out of the simulation box. This setup allowed completely free internal motion of the protein, as well as free translation within a radius of 9.8 nm from the center of mass of the protein (the radius of the protein is about 5.6 nm), which simplified the analysis of the simulations.

### Coarse-grained (CG) Molecular Dynamics simulations

CG models were built using the predicted structure for the open and closed states of Ist2, after energy minimization. The minimized atomistic structures were mapped to a CG representation using the Martinize2 (Kroon et al., 2024) with Martini force field (v3.0) (Souza et al., 2021). DSSP (https://github.com/cmbi/dssp) was used to determine the secondary structure. To stabilize the protein’s tertiary and higher-order structures, we applied a network of Gō potentials between the backbone (BB) beads using the GōMartini3 approach (Souza et al., 2024). Contact maps were generated from http://pomalab.ippt.pan.pl/GoContactMap/ using the OV + rCSU variant, which accounts for both van der Waals (vdW) radii overlap (OV) and repulsive chemical structural units (rCSU), incorporating the chemical properties of the atoms in contact. The depth of the Lennard-Jones potential, in the Gō-like model, were ε = 12.0 kJ/mol and 11.0 kJ/mol in the open and closed states, respectively.

We used the insane.py python script to embed the CG protein into a lipid bilayer (Wassenaar et al., 2015). Systems were solvated using the standard Martini water model and neutralized with 0.15 M KCl. The box dimensions of the system were 20 × 20 × 17 nm^3^. Each system consisted of ∼58,000 CG beads.

CG simulations of Ist2 (open and closed states; Ca^2+^-unbound form) were performed in 2 different lipid mixtures mimicking the ER composition (see table 1), for a total of 4 systems. Control systems of both membranes without Ist2 protein (membrane only system) were also prepared. The details of the simulations performed, with their system composition are summarized in Table 2.

All simulations were carried out with the Gromacs software (Abraham et al., 2015), version 2019. Non-bonded interactions were calculated using the Verlet cutoff scheme, and a buffer tolerance of 0.005 kJ/mol; the interaction cutoff of 1.1 nm. The reaction field approach was used for long-range electrostatic interactions, with a Coulomb cut off of 1.1 nm and a Potential shift Verlet modifier, as implemented in Gromacs.

Energy minimization was carried out using the steepest descent algorithm for 50,000 steps, followed by two 250-ns equilibrations, during which the backbone atoms from the protein structure were positionally restrained (k = 1000 kJ mol^−1^ nm^−2^ and 100 kJ mol^−1^ nm^−2^, respectively). The temperature and pressure were controlled with a velocity-rescale thermostat (Bussi et al., 2007) (reference temperature T = 298 K, coupling constant τ_T_ = 1 ps) and a Parrinello–Rahman semi-isotropic barostat (p = 1 bar, τ_p_ = 12 ps, compressibility β = 3 × 10^−4^ bar^−1^), respectively. Solvent and solute (lipids and protein) beads were coupled separately to the temperature bath. The leap-frog integrator was used, with a time step of 20 fs.

After these two equilibration steps, production runs were carried out in 3 independent replicas, with different initial velocities. Each run was carried out for 20 µs, resulting in a total of 60 µs per system, and 360 µs in total (2 protein systems and one membrane-only as a control). Simulation parameters were as above, except for the absence of position restraints on the protein. Coordinates were saved every 500 ps, resulting in 100,000 snapshots per trajectory for subsequent analysis.

### Simulation analysis

Prior to analysis, trajectories were translationally and rotationally fitted to the protein backbone (BB beads in Martini proteins, see (Monticelli et al., 2008) using Gromacs tools (gmx trjconv). For protein−lipid contact analysis we used the Prolint (Sejdiu and Tieleman, 2021) package. For scrambling analysis, we used the MDanalysis library (Gowers et al., 2016), the LiPyphilic tool (Smith and Lorenz, 2021), and in-house Python scripts. To quantify lipid flip-flop events, we monitored the z-coordinate (direction of the membrane normal) of the PO4 group of each lipid over the entire simulation trajectory. An individual lipid molecule was determined to belong to the upper (lower) leaflet if the z coordinate of its PO4 bead (representing the phosphodiester group in phospholipids) at t = 0 of the production run was higher (lower) than the z coordinate of the membrane center of mass. A complete scrambling event was defined as the movement of a PO4 group from one leaflet to the opposite leaflet, followed by its sustained presence there for a duration of 5 nanoseconds. Statistical uncertainties are reported in tables 1-4 as the standard error on each metric based on 3 independent samples (i.e., the 3 independent simulation replicas obtained for each system).

The bilayer thickness over both leaflets was calculated by means of the g_lomepro tool (Gapsys et al., 2013). The distances between the PO4 beads of lipids located in the upper and lower leaflets with respect to the z-axis were considered as references, using a grid of 100 × 100 points to define their location. Visual Molecular Dynamics (VMD) version 1.9.3 (Humphrey et al., 1996) was used for visual inspection.

### Yeast strain construction and manipulation

Yeast strains are listed in Supplemental Table S2. Yeast were grown in rich medium (YPD) or in synthetic dextrose (SD) medium containing 2% (wt/vol) glucose and appropriate amino acid drop- out mix (MP Biomedicals). Yeast was transformed by standard lithium acetate/polyethylene glycol procedure. SD medium was supplemented with 1 mM ethanolamine when growing *cho1Δ* strain. Deletion and C-ter truncation strains were generated by homologous recombination using appropriate marker (Janke et al., 2004), as indicated. To generate the strains *ist2*ΔN and *ist2-6A* in *ist2*Δ and in *csf1*ΔC *ist2*Δ background, we employed a marker-less CRISPR-Cas9 genomic editing system targeting the *hphMX* antibiotic cassette (Soreanu et al., 2018) inserted at the *IST2* locus, retaining promotor and 3’UTR regions. Hygromycin-sensitive colonies were verified by PCR and sequencing. “Gal-induced” deletion

### Reagents and standards for lipid analysis

Lipid standards (reference, abbreviation) were acquired from Sigma and Avanti Polar Lipids: Egg and 17:1 L-α-lyso-phosphatidylcholine (830071, 855677 LPC), egg or 16:0-18:1 L-α-phosphatidylcholine (840051, 850457 PC), brain or 16:0-18:1 L-α-phosphatidylserine (840032, 840034 PS), liver or 16:0-18:1 L-α-phosphatidylinositol (840042, 850142 PI), heart cardiolipin (840012, CL), egg or 16:0-18:1 L-α-phosphatidic acid (840101, 840857 PA), liver or 16:0-18:1 L-α-phosphatidylethanolamine (840026, 850757 PE), egg L-α-phosphatidylglycerol (841138, PG), N-24:0 (2S-OH) Phytosphingosine C24 (860925, phytoSp), egg ceramide (860051, Cer), 5- cholesten-3β-ol (C8667, Cho), cholest-5-en-3ß-yl heptadecanoate (700186, SE), cholesteryl formate (S448532, ChoF), arachidonic acid (A3611, FFA), 1-oleoyl-rac-glycerol (330724, MG), 1,3-di(cis-9-octadecenoyl)-glycerol (D3627, DG1.3), 1,2-dipalmitoyl-sn-glycerol (800816, DG1.2), 1,2,3-tri-(9Z-octadecenoyl)-glycerol (T7140, TG or TG18:1), 1,2,3-tritetradecanoylglycerol (T5141, TG14:0). Chloroform (CHCl_3_), methanol (MeOH) and hexane (Hex) were HPLC grade; toluene (Sigma, 244511); ethyl acetate (VWR, 23880.290, EtAc). Other reagents: butylhydroxytoluene (Sigma, 34750, BHT), ammonium molybdate tetrahydrate (Sigma, A7302), ascorbic acid (Sigma, 33034), sodium dihydrogen phosphate dihydrate (28011.291), perchloric acid 65-71% (Carlo Erba, 409193), nitric acid 65% (RP Normapur), sulphuric acid 95% (RP Normapur, 20700), phosphoric acid (Sigma, 1.00565), ammonium formate (99%, Acros Organics), copper(II) sulphate pentahydrate (Sigma, C8027).

### Cell growth, lipid extraction and phospholipid quantification

Samples were prepared following the protocol described by (Klose et al., 2012), with a minor modification: cells were cultured in a 1:2.5 liquid-to-air ratio. Briefly, overnight cultures were adjusted to an OD_600_=0.15 in synthetic complete medium using drop-out mix – Ura (MP biochemicals), supplemented with uracil. Cells were harvested by centrifugation in mid-logarithmic phase (OD_600_=0.6), washed once and resuspended in 155 mM ammonium bicarbonate solution. Pellets were snap-frozen in liquid nitrogen, within 12 min from the first centrifugation. Lipid extraction was performed using the MTBE method (Matyash et al., 2008), with modifications. Briefly, 200 µL water was added to samples. After vortexing, samples were transferred into 2 mL plastic tubes with O-ring screw caps containing 0.5 mm glass beads, previously rinsed in MeOH. 300 - 400 µL of MeOH was added to samples and they were homogenized in a Precellys Evolution 2x 20s, 5000 rpm. Then, samples were transferred into glass tubes and the homogenization tubes were rinsed with MeOH by vortexing. MTBE was added to the glass tubes, followed by vortexing at 4°C for 30 min. All extraction solvents contained 50 μg/mL BHT prepared fresh. Extracted samples were kept dried at -20°C under Ar. Phospholipids were quantified using the (Rouser et al., 1970) method, with scaled-down volumes as described in (Dias Araújo et al., 2025).

### Lipid class separation by high performance thin layer chromatography (HPTLC)

For separation of lipid classes, HPTLC instrumentation was used and operated through visionCATS software version 3.0.20196.1. Unless mentioned, instrumentation parameters are as described in (Dias Araújo et al., 2025). PL were dissolved in chloroform as 2 µg/µL and applied (spray 150 nL/s) onto Merck HPTLC glass plates coated with silica gel 60 only (Fig. 4B, 4G) (20x10 cm, layer thickness 200 µm) or embedded with F254 (Fig. 4E) using an automatic TLC sampler 4 (ATS4, CAMAG). Standards were prepared at 1 µg/µL. 2 biological replicates were applied per condition.

To separate polar and non-polar lipids in a single plate, plates were eluted, dried, derivatized and imaged as described in (Dias Araújo et al., 2025). To separate neutral lipid classes only, the plate was washed in CHCl_3_:MeOH (1:1, V:V) in an automated developing chamber (ADC2, CAMAG) up to 85 mm and activated before use (15 min, 100°C). After sample application, the plate was eluted in an automated multiple development chamber (AMD2, CAMAG) with 4 different solvent mixes (V:V) in the following order: 1) CHCl_3_: Hexane: MeOH (45:52:3) up to 55 mm, with MeOH pre- conditioning, 2) Hexane: diethyl ether (80:20) up to 70 mm, 3) Hexane: diethyl ether: acetic acid (80:20:2) up to 72 mm, and 4) Hexane (100%) up to 80 mm. Mix 3 was prepared in one single bottle and used as 100%. Drying time between elutions was 3 min. Images of the 3 plates were treated in Fiji. The areas of the peaks of interest on the chromatograms generated by Fiji were manually delimited and were used to determine the relative lipid composition per lane.

### Live yeast cell imaging

Yeast cells were grown for 14–18 h in appropriate SD medium at 30°C, except for *sec23-1* where cells were grown at 25°C. Cells were diluted and harvested by centrifugation in mid-logarithmic phase (OD_600_=0.6–0.8) and prepared for visualization on glass slides. For LD counting, cells were grown to early stationary phase (1.8>OD_600_>1) and for the pKE41 probe localization, cells were grown for 24h prior to observation. When indicated, LD staining was obtained by incubating cells with 1 mg/mL of BODIPY® 493/503 (ThermoFisher) or with 0.1 mM of AUTODOT™ (Cliniscience) for 30 min. For rapamycin-induced autophagy, cells were grown to early exponential phase (OD_600_=0.3) before being treated with 200 nM of rapamycin for 3h (Liu et al., 2024). Imaging was conducted at room temperature, using a Spinning disk SR system (Olympus), equipped with an oil immersion plan apochromat 60x objective NA 1.42, an sCMOS Fusion BT camera (Hamamatsu) and a spinning-disk confocal system CSU-W1 (Yokogawa). For LD counting experiments and GFP-tagged versions of Ist2, the SoRa spinning-disk module was employed. AUTODOT, BODIPY/GFP-tagged proteins and mScarlet-tagged proteins were visualized with DAPI (447/50), GFP (525/50), and mCherry (593/40) filters, respectively. Cells were imaged in 3 to 15 *z*-sections separated by 0.25 µm. Images were acquired using CellSens software (Olympus) and processed with Fiji (ImageJ).

### Image analysis

Images were processed with Fiji (ImageJ). For the rapamycin-induced autophagy experiment, the number of foci was manually counted from a stack of 3–5 z-sections. LD counting and estimation of LD fluorescence-positive cells using the Plin4-AH probe were performed manually from a stack of 15 z-sections. Steady-state distribution of C2_Lact_–GFP was analyzed from a stack of 3 *z*-section of each cell. The quantification of peripheral peaks (mostly PM and cortical ER signal) and internal (mostly perinuclear ER) was performed by profiling cell signal intensity across a transversal line drawn in the cell with an automated macro. Intensities of the peaks were quantified and normalized relative to the total signal as previously described (D’Ambrosio et al., 2019, 2020).

### Viability and kinetic assays

Yeast cells were grown for 14–18 h at 30°C in appropriate medium (YPD or SD medium), except for *sec23-1*, which was cultured at 25°C. For drop-test assays, cells were adjusted to the lowest OD_600_, serially diluted, and plated onto the appropriate agar media. Plates were incubated for 2-6 days at designated temperatures. For kinetic assays, cells were adjusted to an OD_600_=0.1 in the appropriate SD medium, and grown in technical and in biological duplicates, in 96-well microplates using the Spark^®^ microplate reader (Tecan). For the *ist2Δpsd1Δ* strain, cells were supplemented with 2mM of ethanolamine overnight before the kinetic, in presence or absence of ethanolamine. For each kinetic cycle, with an interval of 10min, the cultures were shaken for 9 min with an amplitude of 2mm and a frequency of 150 rpm, followed by a 1-min settling period before measuring the absorbance at 600nm. Kinetic measurements continued until the controls reached the plateau of the stationary phase.

### Western blot analysis of total yeast extracts

Proteins were extracted from one OD_600_ equivalent of yeast cells grown to mid-logarithmic phase in appropriate SD medium. Protein samples were heated at 55°C for 10 min and 30 µL were loaded on SDS–PAGE (4–20% Mini-PROTEAN TGX Stain-Free, Bio-Rad for Supp Fig. 4C and 10% homemade gel for Supp Fig. 4D) and analyzed by western blotting. Briefly, nitrocellulose membranes were saturated with 5% of milk, incubated ON at 4°C with primary antibody (1:1000, anti-Halo, Promega G9211, and anti-RFP, Chromotek 6G6), washed with TBS-Tween20, incubated 1h at RT with secondary antibody (Invitrogen^TM^ AlexaFluor^TM^ anti-mouse 568, or anti-rabbit 488 respectively), washed with TBS-Tween20 and revealed with an Odyssey imaging system.

### Statistical analysis

For cross-analysis a Shapiro test was used to control the normality of the values and a Bartlett test for the homogeneity of variances. Kruskal-Wallis test followed by Dunn’s post-hoc test was then used to test the equality of ranks across conditions. The representation of p values is as follows: ns, non-significant *<0.05, **<0.01, ***<0.001, ****<0.0001. The nomenclature (N=x) refers to the number of biological replicates and (n=x) refers to the number of measurements per replicate.

## Notes

### Competing Interest Statement

The authors have declared no competing interest.

