## Supplemental data for "Ist2, a protein involved in phosphatidylserine transport, is an ER lipid scramblase"

Sebinelli, Syska et al., Supplemental Materials

Supplemental Figures S1-S3

Supplemental Tables 1-4

Supplemental Movie 1

### Supplemental Figure 1

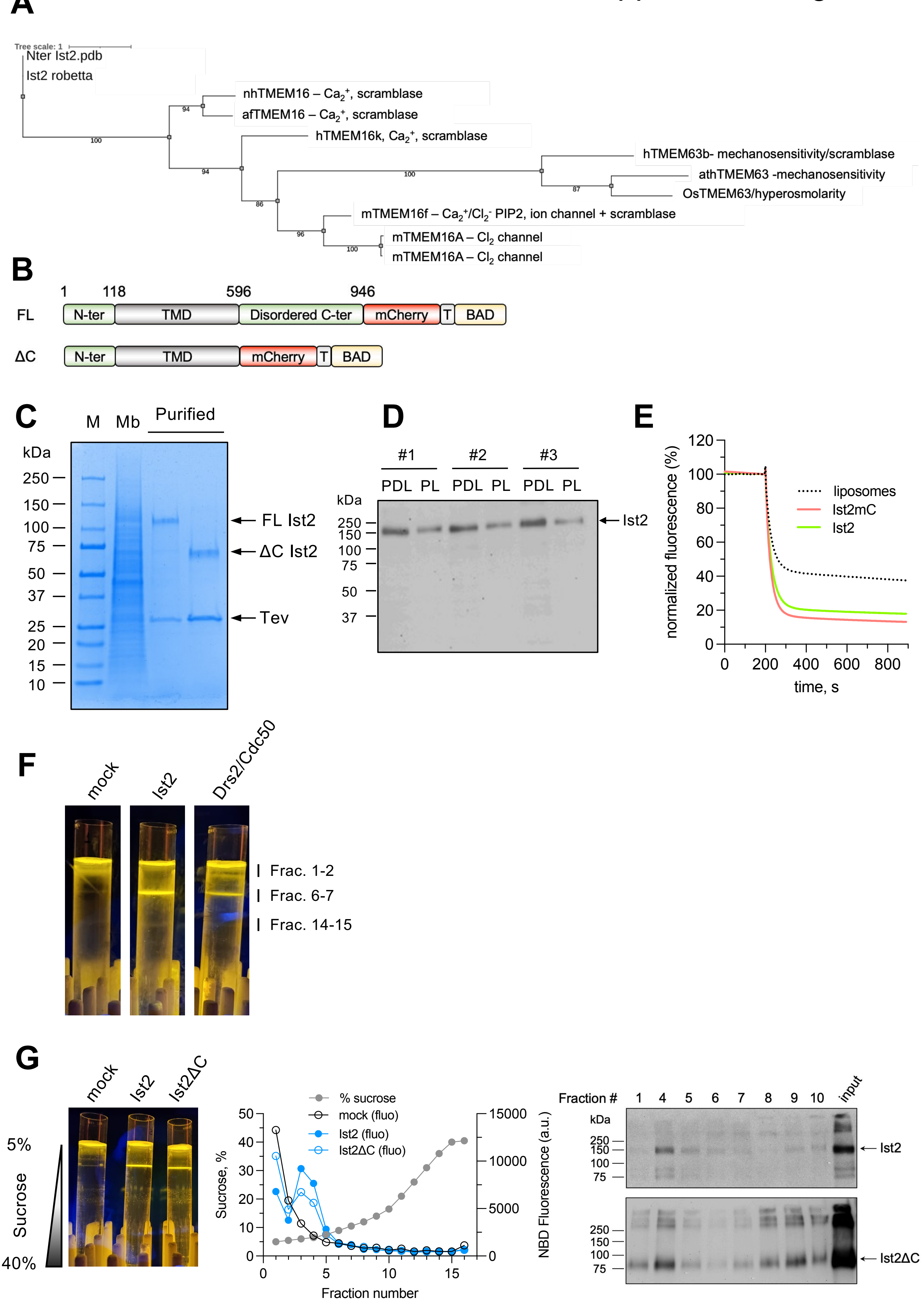

**Supplemental figure 1. (A)** Phylogenetic tree of the top homologous proteins to Ist2 identified by I-TASSER. Crystal structures of nhTMEM16 (4WIS), afTMEM16 (7RWJ), hTMEM16K (5OC9), hTMEM63B (8EHX), athTMEM63 (8GRO), Os/hyperosmolarity (6OCE), mTMEM16F (8TAI), and mTMEM16A (5NL2, 7ZK3), along with predicted models of Ist2 from I-TASSER and Robetta, were aligned using PROMALS3D. The alignment was then analyzed using IQ-TREE (maximum likelihood algorithm), and the resulting phylogenetic tree was visualized with iTOL. **(B)** Schematic of the constructs used for the purification and reconstitution of Ist2. The C-terminus of Ist2 was fused to a biotin-acceptor domain (BAD) for in vivo biotinylation in yeast and affinity purification on streptavidin beads, and to mCherry, for screening of solubilization conditions by Fluorescence Size-Exclusion Chromatography. A Tobacco Etch Virus (TEV) protease cleavage site was added between the BAD and mCherry to release Ist2-mCherry from streptavidin beads. A construct lacking the C-terminus (ΔC), predicted to be disordered, was included. **(C)** SDS-PAGE analysis of full-length and Ist2-ΔC purification on streptavidin beads. Crude yeast membranes (Mb) (of which Ist2 represents 2-3 %) and proteins recovered upon TEV protease cleavage on streptavidin beads ($E_{strep}$) were loaded on the gel and visualized by Coomassie Blue staining. M, molecular weight marker. **(D)** Reconstitution efficiency of Ist2 following treatment of protein/detergent/lipid (PDL) extracts with biobeads, as assessed by western-blot analysis of PDL extracts and the resulting proteoliposomes (PL). Ist2 was detected using an antibody against mCherry. **(E)** Picture of the sucrose gradient after ultracentrifugation for 20 h at 36,000 rpm. C12-NBD-PS fluorescence was revealed using a fluorescence imager. **(F)** NBD-PS quenching of liposomes or proteoliposomes reconstituted with either mCherry-tagged or untagged Ist2. The initial fluorescence was recorded for 200 s before dithionite addition. **(G)** Sucrose gradient fractionation of reconstituted full-length (Ist2) and C-terminally truncated Ist2 (Ist2-ΔC). Left panel: picture of the sucrose gradient after ultracentrifugation for 20 h at 36,000 rpm. Fractions collected from the top of the gradient upon ultracentrifugation were probed for sucrose concentration and NBD fluorescence (middle panel) and the presence of Ist2 was detected using an anti-mCherry antibody (right panel).

### Supplemental Figure S2

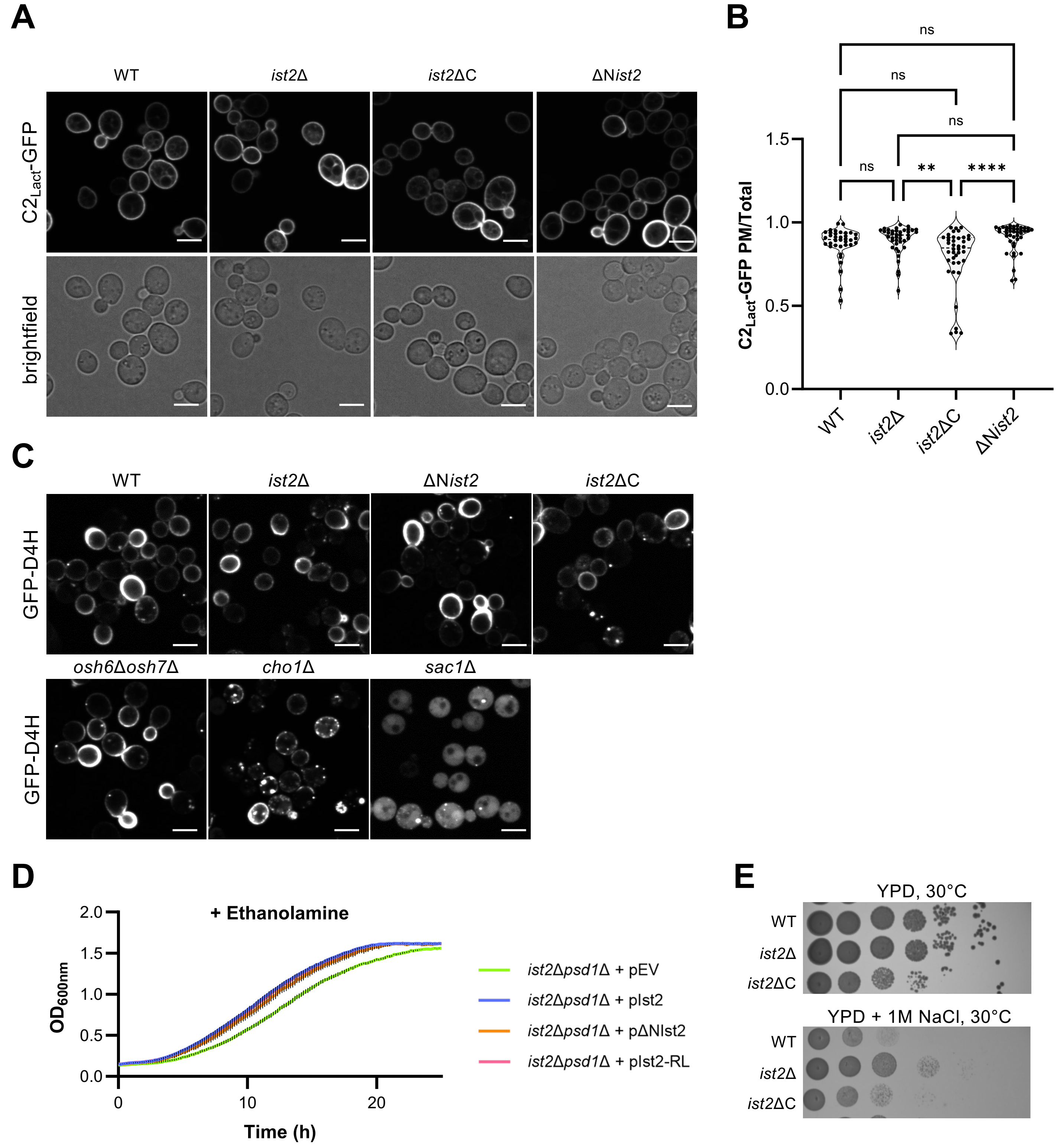


**Supplemental figure 2. Yeast data related to Figure 4. (A)** Steady-state distribution of PS in indicated yeast cells. Corresponding fluorescent microscopy images show C2$_{Lact}$-GFP in WT cells and cells with chromosomal mutations in *IST2*, as indicated. Scale bar = 5 µm. **(B)** Quantification of relative C2$_{Lact}$-GFP peak intensity at PM, normalized to total cellular fluorescence (PM and internal peaks). Each violin plot represents the quantification of 40 individual cells. Statistical significance was determined using the Kruskal-Wallis test followed by Dunn's post-hoc test, P>0.05 (ns), P<0.01 (**), P<0.0001 (****). The graph represents data from a representative of 3 experiments. **(C)** Steady-state distribution of sterols in indicated yeast cells, as assessed by the sterol probe GFP-D4H. Scale bar = 5 µm. **(D)** Growth kinetics of *ist2Δpsd1Δ* cells expressing WT and mutated versions of Ist2, tagged with BFP at N-ter, grown in SC–His medium at 30°C in the presence of ethanolamine. **(E)** Growth of indicates strains on YPD, supplemented or not with 1M of NaCl. Ten-fold serial dilutions of overnight cultures were spotted on the plates and incubated for 3 days at 30ºC.

Supplemental Figure S3

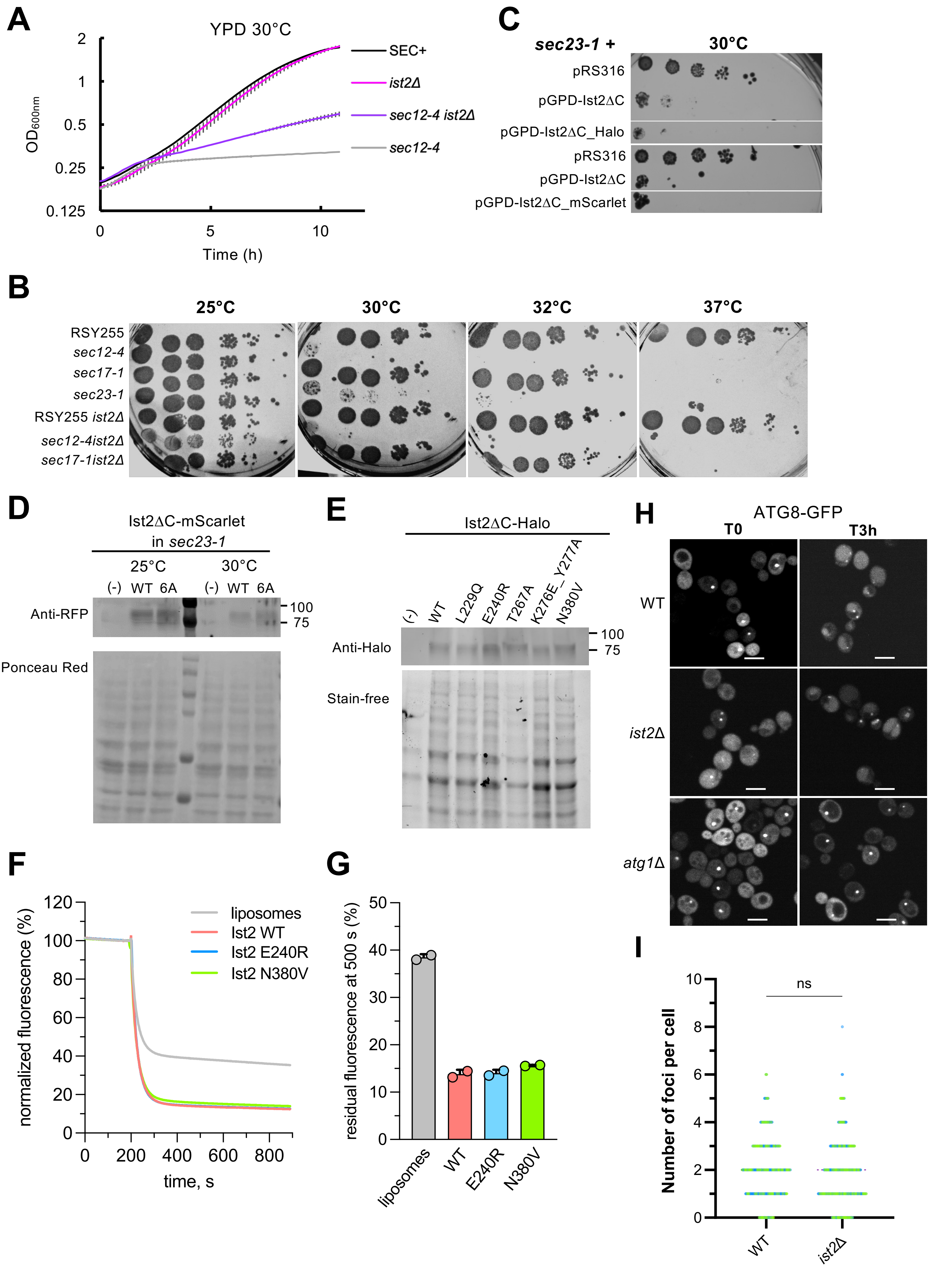

**Supplemental figure 3. Yeast and in vitro data data related to Fig. 5. (A)** Growth kinetics of indicated yeast strains, grown in YPD at 30°C. The growth curves represent the mean ± range of absorbance at 600 nm ($OD_{600}$) of two independent biological replicates, each measured in technical duplicates over time (hour). **(B)** Viability assay of the indicated yeast strains grown on YPD agar. Ten-fold serial dilutions of overnight cultures were spotted onto the plates and incubated for 2 days at 25°C, 30°C, 32°C and 37°C. **(C)** Viability assay of *sec23-1* cells overexpressing Ist2ΔC, tagged or untagged, from a centromeric plasmid with a GPD promotor. Ten-fold serial dilutions of overnight cultures were spotted onto SC-URA plates and incubated for 6 days at 30°C. **(D)** Expression of Ist2ΔC-mScarlet and Ist2Δ$C_{6A}$-mScarlet in *sec23-1* cells at 25°C and 30°C. Anti-RFP western blot detects the versions of Ist2ΔC-mScarlet at the expected size (75kDa). Ponceau red staining shows the total protein transfer to the nitrocellulose membrane following 10% SDS-PAGE gel. (-) refers to empty plasmid control. **(E)** Expression of Ist2ΔC-Halo WT and mutated versions in *sec23-1* cells at 30°C. Anti-Halo Western Blot detects Ist2ΔC-Halo WT and variants. Stain-free imaging shows the total protein loading onto a 4-20% gradient SDS-PAGE gel. (-) refers to empty plasmid control. **(E)** Single point mutations E240R and N380V do not affect NBD-PS scrambling of Ist2. Traces represent the mean of 2 transport experiments from independent reconstitutions. **(F)** Quantification of residual fluorescence in liposomes versus proteoliposomes upon dithionite quenching. The residual fluorescence measured in panel E at 500 s was plotted. Data are a mean ± s.d. of 2 biological replicates (independent reconstitutions). **(H)** Distribution of ATG8-GFP in WT, *ist2*Δ and *atg1Δ* cells before and after 3h of rapamycin-induced autophagy. Scale bar = 5 µm. **(I)** Quantification of the number of foci per cell in WT and *ist2*Δ cells 3h after rapamycin-induced addition (n≥100, N=2). The significance was assessed by Mann-Whitney test, P>0.05 (ns).

### Supplemental Tables

**Supplemental Table 1: Plasmids used in this study.** To note that all plasmids carry an ampicillin antibiotic resistance. For truncated versions, the retained codon positions are indicated in brackets.

| Plasmids | Alias | Description | References |
|---|---|---|---|
| **pJET1.2** | pJET1.2 | Cloning vector, AmpR | Thermo Scientific™ |
| **pJET-Ist2** | pJET-Ist2 | pJET1.2 derivative, containing *IST2* gene | This work |
| **pYeDP60** | pYeDP60 | *2µ, URA3/ADE2,* GALpr- TEV-BAD | Pompon et al, 1996 |
| **pYeDP60-Ist2-Tev-BAD** | | pYeDP60-based, GALpr-Ist2-TEV-BAD | This work |
| **pGPDmcherryp413** | | pRS413-based (*CEN, HIS3*), GPDpr-mCherry | Pr Sophie Dupré (UVSQ) |
| **pYeDP60-Ist2_mcherry-Tev-BAD** | | pYeDP60-based, GALpr-Ist2_mCherry-TEV-BAD | This work |
| **pYeDP60-Ist2_E240R-mcherry-Tev-BAD** | | pYeDP60-based, GALpr-Ist2_E240R_mCherry-TEV-BAD | This work |
| **pYeDP60-Ist2_N380V-mcherry-Tev-BAD** | | pYeDP60-based, GALpr-Ist2_N380V_mCherry-TEV-BAD | This work |
| **pYeDP60-Ist2ΔC -Tev-BAD** | | pYeDP60-based, GALpr-Ist2[1-600]-TEV-BAD | This work |
| **pYeDP60-Ist2ΔC-mcherry-Tev-BAD** | | pYeDP60-based, GALpr-Ist2[1-600]_mCherry-TEV-BAD | This work |
| **pRS413** | pRS413/pEV | *CEN, HIS3* | Lab collection |
| **pJMD_07** | pIst2 | pUG34-based (*CEN, HIS3*), Ist2pr-BFP-Ist2 | D'Ambrosio et al., 2020 |
| **pJMD_09** | pΔN-Ist2 | pUG34-based (*CEN, HIS3*), Ist2pr-BFP-ΔN-Ist2 [590-946] | D'Ambrosio et al., 2020 |
| **pJMD_21** | pIst2-RL | pUG34-based (*CEN, HIS3*), Ist2pr-BFP-Ist2-RL [595-946 randomized linker], *HIS3, CEN* | D'Ambrosio et al., 2020 |
| **pYM-N23** | | *NAT*, GALpr, template for gene expression | Janke et al., 2004 |
| **pYM-N23_GAL-CHO1::NAT** | | pYM-N23-based, GALpr-*CHO1*::*NAT* | This work |
| **LactC2-GFP** | pC2lact-GFP | GPDprC2Lact-GFP, *CEN*, *URA3,* Addgene plasmid # 22852 | Yeung et al., 2008 |
| **D4H GFP ura** | pGFP-D4H | CUP1pr-GFP-D4H insertion in pRS316, *CEN, URA3* | Del Dedo et al., 2021 |
| **pRS316** | pRS316/p0 | *CEN, URA3* | Lab collection |
| **pRS416** | pADH | pRS416-based, ADH1pr | Lab collection |
| **pVA_72** | pADH-Ist2ΔC | ADH1pr-Ist2ΔC, Ist2 [1-600], pRS416, *URA3* | This work |
| **pRS416** | pGPD | pRS416-based, GPDpr | Lab collection |
| **pVA_73** | pGPD-Ist2ΔC | GPDpr-Ist2ΔC, Ist2 [1-600], pRS416, *URA3* | This work |
| **pCS_35** | pGPD-Ist2ΔC_6A | Ist2[1-600] *L229A, E240A, T267A, R359A, Q363A, Y366A;* Site-directed mutagenesis of pVA73 | This work |
| **pCS_20** | pGPD-Ist2ΔC_E240R | Ist2[1-600] *E240R,* Site-directed mutagenesis of pVA_73 | This work |
| **pCS_13** | pGPD-Ist2 | pRS416-based, GPDpr-Ist2 (full length) | This work |
| **pCS_33** | pGPD-Ist2_6A | Ist2 *L229A, E240A, T267A, R359A, Q363A, Y366A;* Site-directed mutagenesis of pCS_13 | This work |
| **pCS_27** | pGPD-Ist2_E240R | Ist2 *E240R,* Site-directed mutagenesis of pCS_13 | This work |
| **pBBK83** | ERG6-Link-Scarlet | pHomL-RPL18Bp-ERG6-Link-Scarlet(rfp)-SSA1t-HomR (AmpR), Addgene 179067 | Bean et al., 2022 |
| **pCS_14** | pGPD-Ist2ΔC_mScarlet | GPDpr-Ist2ΔC_mScarlet, insertion of mScarlet tag in pVA_73 | This work |
| **pCS_36** | pGPD-Ist2ΔC_mScarlet_6A | Ist2[1-600] *L229A, E240A, T267A, R359A, Q363A, Y366A*-mScarlet*;* Site-directed mutagenesis of pCS_14 | This work |
| **pBS-SKII-3XHA-HALO-NAT** | pHALO-NAT | pBS-SKII-3XHA-HALO-NAT (AmpR), Addgene 188931 | Bean et al., 2022 |
| **pCS_07** | pGPD-Ist2ΔC_Halo | GPDpr-Ist2ΔC_Halo, insertion of Halo tag in pCS_07 | This work |
| **pCS_11** | pGPD-Ist2ΔC_Halo_L229Q | Ist2[1-600] *L229Q*-Halo*,* Site-directed mutagenesis of pCS_07 | This work |
| **pCS_09** | pGPD-Ist2ΔC_Halo_E240R | Ist2[1-600] *E240R*-Halo*,* Site-directed mutagenesis of pCS_07 | This work |
| **pCS_08** | pGPD-Ist2ΔC_Halo_T267A | Ist2[1-600] *T267A*-Halo*,* Site-directed mutagenesis of pCS_07 | This work |
| **pCS_12** | pGPD-Ist2ΔC_Halo_K276E_Y277A | Ist2[1-600] *K276E Y277A*-Halo, Site-directed mutagenesis of pCS_07 | This work |
| **pCS_10** | pGPD-Ist2ΔC_Halo_N380V | Ist2[1-600] *N380V*-Halo, Site-directed mutagenesis of pCS_07 | This work |
| **pGFP-ATG8_LEU** | | pRS315-based (*CEN, LEU2*), ADHprGFP-ATG8 | |
| **pADH-P4m12nt-GFP** | pKE41 | pRS416-based, ADH1pr-Plin4m12nt (246-377) 2K->Q, 2T->V | Čopič et al., 2018 |
| **pVA_469** | pCY5 (HygrMX) | CASpr (*2µ, URA3*), gRNA-*HghMX* | Soreanu et al., 2018 |

| ***S.cerevisiae* S288 Alias** | **Strains** | **Background** | **Genotype** | **References** |
|---|---|---|---|---|
| **MBY03** | GFP::IST2 | BY4742 | *MATα his3Δ1 leu2Δ0 lys2Δ0 ura3Δ0 GFP-IST2* | This work |
| **ACY415** | *ist2*Δ$C_{590}$::GFP | BY4741 | *MATa his3Δ1 leu2Δ0 met15Δ0 ura3Δ0 ist2Δ590-946::GFP::HygrMX* | This work |
| **MBY05** | GFP::*ist2*Δ$N_{490}$ | BY4742 | *MATα his3Δ1 leu2Δ0 lys2Δ0 ura3Δ0 GFP::ist2Δ*1-490 | This work |
| **BY4742** | WT : BY4742 | BY4742 | *MATα his3Δ1 leu2Δ0 lys2Δ0 ura3Δ0* | Euroscarf |
| **VAY2784** | *ist2Δ* | BY4742 | *MATα his3Δ1 leu2Δ0 lys2Δ0 ura3Δ0 ist2Δ::HygMX* | D'Ambrosio et al., 2020 |
| **CSY01** | *ist2ΔN* | BY4742 | *MATα his3Δ1 leu2Δ0 lys2Δ0 ura3Δ0 ist2Δ1-490* | This work |
| **CSY05** | *ist2-6A* | BY4742 | *MATα his3Δ1 leu2Δ0 lys2Δ0 ura3Δ0 ist2 L229A, E240A, T267A, R359A, Q363A, Y366A* | This work |
| **BYcho1** | *cho1Δ* | BY4742 | *MATα his3Δ1 leu2Δ0 lys2Δ0 ura3Δ0 cho1Δ::His3MX* | D'Ambrosio et al., 2020 |
| **SGAY1635** | *osh6Δosh7Δ* | SGA Y7039 | *MATα can1Δ::STE2pr-LEU2 lyp1Δ ura3Δ0 leu2Δ0 his3Δ1 met15Δ0 osh6Δ::HygMX osh7Δ::NatMX* | Maeda et al., 2013 |
| **MdPY11** | *psd1D::kanMX ist2D::natMX* | BY4741 | *MATa his3Δ1 leu2Δ0 met15Δ0 ura3Δ0* | This work |
| **BY4741** | WT : BY4741 | BY4741 | *MATa his3Δ1 leu2Δ0 met15Δ0 ura3Δ0* | Euroscarf |
| **MdPY13** | *ist2ΔC* | BY4741 | *MATa his3Δ1 leu2Δ0 met15Δ0 ura3Δ0 ist2Δ590-946::HygMX* | D'Ambrosio et al., 2020 |
| **ACY / VAY** | BY4741 Gal-*CHO1::NAT* | BY4741 | *MATa his3Δ1 leu2Δ0 met15Δ0 ura3Δ0* BY4741 Gal-*CHO1::NAT* | This work |
| **ACY / VAY** | *ist2Δ* Gal-*CHO1::NAT* | BY4742 | *MATα his3Δ1 leu2Δ0 lys2Δ0 ura3Δ0 ist2Δ::HygMX* Gal-*CHO1::NAT* | This work |
| **ACY / VAY** | *ist2Δ*N Gal-*CHO1::NAT* | BY4742 | *MATα his3Δ1 leu2Δ0 lys2Δ0 ura3Δ0 ist2Δ1-490* Gal-*CHO1::NAT* | This work |
| **RSY255** | RSY255 | RSY255 | *MATα ura3-52 leu2,3,-112* | Schekman Lab |
| **ACY429** | *ist2Δ* RSY255 | SEC+ | *MATα ura3-52 leu2,3,-112 ist2Δ::KanMX* | This work |
| **RSY263** | *sec12-4* | SEC+ | *MATα ura3-52 his4-619 sec12-4* | Schekman Lab |
| **ACY430** | *sec12-4 ist2Δ* | SEC+ | *MATα ura3-52 his4-619 sec12-4 ist2Δ::KanMX* | This work |
| **RSY281** | *sec23-1* | SEC+ | *MATα ura3-52 his4-619 sec23-1* | Schekman Lab |
| **ACY418** | *csf1ΔC* | BY4741 | *MATa his3Δ1 leu2Δ0 met15Δ0 ura3Δ0 csf1Δ797-2958::NatMX* | This work |
| **ACY419** | *ist2Δ csf1ΔC* | BY4742 | *MATα his3Δ1 leu2Δ0 lys2Δ0 ura3Δ0 ist2Δ::HYGMX csf1Δ797-2958::NatMX* | This work |
| **ACY427** | *ist2ΔC csf1ΔC* | BY4741 | *MATa his3Δ1 leu2D0 met15Δ0 ura3Δ0 ist2Δ600-946::HygMX csf1Δ797-2958::NatMX* | This work |
| **CSY06** | *ist2ΔN csf1ΔC* | BY4742 | *MATα his3Δ1 leu2Δ0 lys2Δ0 ura3Δ0 ist2Δ1-490 csf1Δ797-2958::NatMX* | This work |
| **CSY07** | *ist2-6A csf1ΔC* | BY4742 | *MATα his3Δ1 leu2Δ0 lys2Δ0 ura3Δ0 ist2 L229A, E240A, T267A, R359A, Q363A, Y366A csf1Δ797-2958::NatMX* | This work |
| **MFY62** | *sac1Δ* | SEY6210.1 | *MATα ura3-52 leu2-3,112 his3Δ200 trp1-Δ901 lys2-801 suc2Δ9 sac1Δ::TRP1* | Emr lab (Foti et al., 2001) |
| ***atg1Δ*** | *atg1Δ::KanMX* | BY4741 | *MATa his3Δ1 leu2Δ0 met15Δ0 ura3Δ0* | Euroscarf |

**Supplemental Table 2: Yeast strains used in this study.** Numbers after the Δ symbol indicate the starting position or the codon range of the deleted region.

| Lipid type | | Unsaturation (sn1-sn2) | Simple lipid mixture | Percentage | Complex lipid mixture | Percentage |
|---|---|---|---|---|---|---|
| PC | DYPC | 16:1-16:1 | 300 | 50% | 120 | 20 % |
| | YOPC | 18:1-16:1 | - | - | 120 | 20 % |
| PE | DYPE | 16:1-16:1 | 150 | 25% | 60 | 10 % |
| | YOPE | 18:1-16:1 | - | - | 60 | 10 % |
| PS | DYPS | 16:1-16:1 | 150 | 25% | - | - |
| | YOPS | 18:1-16:1 | - | - | 60 | 10 % |
| PI | PYPI | 16:1-16:0 | - | - | 60 | 10 % |
| | POPI | 18:1-16:0 | - | - | 60 | 10 % |
| PA | YOPA | 18:1-16:1 | - | - | 30 | 5 % |
| Ergosterol | (ERGO) | - | - | - | 30 | 5 % |
| lipids per leaflet | | | 600 | - | 600 | - |

**Supplemental Table 3. Lipid composition in AA simulations.** We used two different lipid mixtures mimicking the ER composition, here named "simple lipid mixture" and "complex lipid mixture". In CG simulations, we used the same mixtures, replacing PYPI with POPI (they are identical in the Martini CG force field) and YOP* with DOP*.

| Systems | Membrane (mixture) | Resolution | Atoms, Particles | Equilibration time (ns) | Production time (µs) |
|---|---|---|---|---|---|
| Open state-$Ca^{2+}$ | Simple | AA | 707448 | 2×250 | - |
| Open state-$Ca^{2+}$ | Complex | AA | 696813 | 2×250 | - |
| Open state | Simple | AA | 707353 | 2×250 | 2×1 |
| Closed state | Simple | AA | 697564 | 2×250 | 2×1 |
| Open state | Simple | CG | 58308 | 2×250 | 3×20 |
| Open state | Complex | CG | 57847 | 2×250 | 3×20 |
| Closed state | Simple | CG | 58735 | 2×250 | 3×20 |
| Closed state | Complex | CG | 58080 | 2×250 | 3×20 |
| Membrane-only | Simple | CG | 21871 | 1×250 | 3×20 |
| Membrane-only | Complex | CG | 21292 | 1×250 | 3×20 |

**Supplemental Table 4. List of AA and CG MD simulations carried out.**
